# Bioelectrical Interfaces with Cortical Spheroids in Three-Dimensions

**DOI:** 10.1101/2020.11.29.401638

**Authors:** Anna Kalmykov, Jay W. Reddy, Esther Bedoyan, Yingqiao Wang, Raghav Garg, Sahil K. Rastogi, Devora Cohen-Karni, Maysamreza Chamanzar, Tzahi Cohen-Karni

## Abstract

Three-dimensional (3D) neuronal spheroid culture serves as a powerful model system for the investigation of neurological disorders and drug discovery. The success of such a model system requires techniques that enable high-resolution functional readout across the entire spheroid. Conventional microelectrode arrays and implantable neural probes cannot monitor the electrophysiology activity across the entire native 3D geometry of the cellular construct. Here, we demonstrate a 3D self-rolled biosensor array (3D-SR-BA) integrated with a 3D cortical spheroid culture for simultaneous *in-vitro* electrophysiology recording, functional Ca^2+^ imaging, and drug effect monitoring. We have also developed a signal processing pipeline to detect neural firings with high spatiotemporal resolution from the electrophysiology recordings based on established spike sorting methods. The 3D-SR-BAs cortical spheroid interface provides a stable, high sensitivity recording of neural action potentials (< 50 μV peak-to-peak amplitude). The 3D-SR-BA is demonstrated as a potential drug screening platform through the investigation of the neural response to the excitatory neurotransmitter glutamate. Upon addition of glutamate, the neuronal firing rates increased notably corresponding well with the functional Ca^2+^ imaging. Our entire system, including the 3D-SR-BA integrated with neural spheroid culture, enables simultaneous electrophysiology recording and functional Ca^2+^ imaging with high spatiotemporal resolution in conjunction with chemical stimulation. We demonstrate a powerful toolset for future studies of tissue development, disease progression, and drug testing and screening, especially when combined with native spheroid cultures directly extracted from humans.

## Introduction

Three dimensional (3D) cell culture closely mimics the physiology and development of native tissue and serves as a powerful system for disease modeling and designing new therapeutics.^1–7^ Spheroids, the most common 3D tissue model, are based on the cells’ tendency to aggregate, forgoing a scaffold or extracellular matrix and thus reducing the batch-to-batch variation.^8^ 3D cultures of neuronally differentiated stem cells have enabled *in vitro* modeling of diseases such as autism,^3^ microcephaly,^6^ and Alzheimer’s disease.^4^ ^3, 9^ The success of these model systems requires techniques to probe the rich structure and function of the 3D cellular construct with high-resolution functional readout.

3D cell culture holds a great promise in personalized medicine,^1–6^ as it bridges the gap between 2D cell culture and *in vivo* testing. For example, 2D *in vitro* models of the central nervous system (CNS), do not recapitulate axonal regeneration, while in 3D models, synaptic connections and myelinated axons are formed.^10^ Therefore, proper neurodegenerative disease modeling necessitates 3D tissue models.^2, 4–5, 10^ Using human derived *in vivo*-like 3D tissue models may reduce animal testing in drug toxicology studies.^8^ For neural constructs, functionality is most accurately characterized by recording neuronal electrophysiology (ephys).^10^ Recording the electrical activity of neural spheroids in their native 3D architecture over time with high spatial and temporal resolution is critical for developing disease models and validating potential treatments.^10^

3D neural structures have been investigated by conventional neural recording techniques. Patch clamp is limited to recordings from sectioned organoids, and from a few individual cells.^9^ High-density implantable neural probes have been used to record neural activity in the probe vicinity.^11^ Two-dimensional (2D) microelectrode arrays (MEAs) have been used to interface with neuronal organoids but exhibit a limited 2D interface with the organoid.^12–13^ Optical techniques, such as functional Ca^2+^ imaging, which provide high spatial resolution and throughput, lack the temporal precision of ephys techniques (msec resolution), required to study the temporal dynamics of neural activity.^14^

Here, we present a platform consisting of a 3D self-rolled biosensor array (3D-SR-BA)^15^ that wraps around the 3D structure of a rat cortical spheroid. We demonstrate a platform for studying 3D neural spheroids, including cell culture, 3D self-rolling electrode arrays (Figure 1 A I, II), chemical neuronal stimulation, simultaneous ephys recording, and functional Ca^2+^ imaging. We have developed a complete signal processing pipeline to analyze ephys recordings in conjunction with functional Ca^2+^ imaging of the neural activity across different regions of the spheroid in response to glutamate stimulation. The pipeline includes identification of localized neural activity and isolation of single units using spike sorting analysis to obtain firing rates of individual neurons (Figure 1 A III, IV). The demonstrated results in this paper exemplify the utility of this powerful platform for studying the effect of chemical stimulation (i.e., glutamate) on the neuronal response in spheroids with high spatiotemporal resolution using distributed ephys recording and high throughput Ca^2+^ imaging. Ephys recording using 3D-SR-BA will allow unprecedented access to the 3D structure of the organoid for recording neuronal activity with high temporal resolution, while functional Ca^2+^ imaging provides a complementary readout mechanism to capture the holistic activity of the tissue with high spatial resolution within the field of view. This novel platform provides tissue accessibility and high throughput long-term 3D neuronal recording, enabling exciting future biological applications, such as drug screening for neurological disorders.

**Figure 1.**
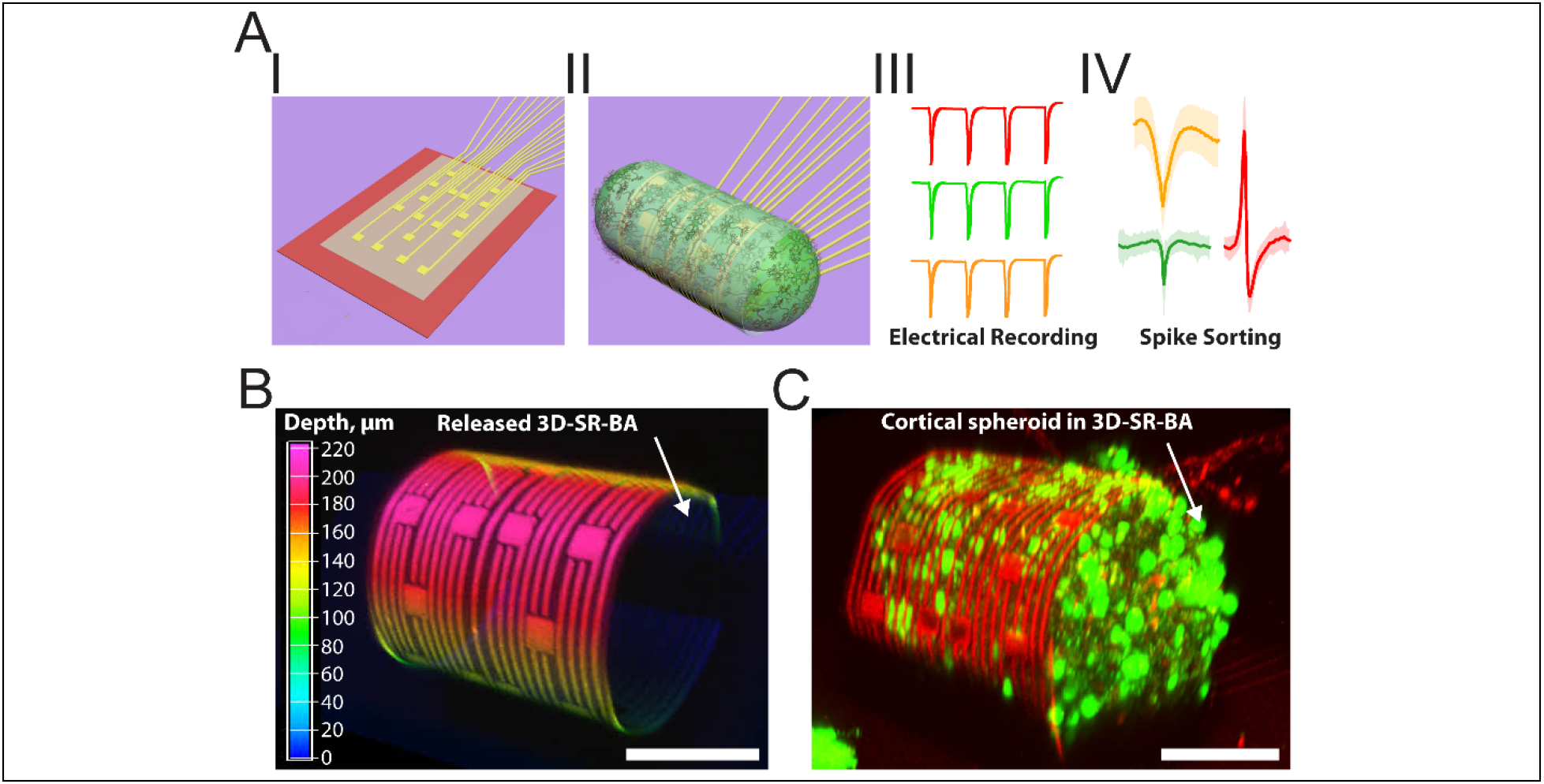
Three-dimensional (3D) self-rolled biosensor array (3D-SR-BA) for 3D recording from cortical spheroids. **(A)** A schematic of the experiment: (I) The 3D-SR-BA is released to conform in 3D, (II) A cortical spheroid is encapsulated in the 3D-SR-BAs, and (III) The electrical recording is obtained. (IV) The recorded data is processed, and spike-sorting algorithms are used to identify individual neuronal activity. **(B)** A 3D confocal microscopy image of 3D-SR-BA. Color bar represents the depth in μm. **(C)** A 3D confocal microscopy image of cortical spheroid labeled with Ca^2+^ indicator dye (Cal-520, green fluorescence) encapsulated by the 3D-SR-BA. Scale bars are 100 μm.

## Materials and Methods

### 3D sensor array fabrication

The 3D-SR-BAs were fabricated following a previously published microfabrication process, outlined in Figure S1.^15–17^ Si substrates with a 600 nm wet thermal oxide (p-type, ≤0.005 Ω cm, Nova Electronic Materials Ltd., catalog no. CP02 11208-OX) were cleaned with acetone and isopropyl alcohol (IPA) in an ultrasonic bath for 5 min each, N2 blow dried, and treated with O2 plasma at 100 W power for 1 min (Barrel Asher, International Plasma Corporation). The substrate was coated with 300 nm lift-off resist (LOR3A, Kayaku Advanced Materials) and 500 nm positive photoresist (Shipley S1805, Kayaku Advanced Materials). Outer electrode interconnects were defined by ultraviolet (UV) lithography using a mask aligner (MA6, SUSS MicroTec SE) followed by development for 1 min (CD26 developer, Kayaku Advanced Materials). 10 nm Cr (99.99%, R.D. Mathis Co.) and 75 nm Au (99.999%, Praxair) were deposited using thermal evaporation (Covap II, Angstrom Engineering Inc) as the interconnect metals. The LOR3A/Shipley 1805/metal stack was lifted off using Remover PG (Kayaku Advanced Materials) to pattern the electrode interconnects. The wafer was washed with acetone and IPA, N2 blow-dried, and treated with O2 plasma at 50 W power for 1 min. A 200 nm Ge (99.999%, Kurt J. Lesker Company) sacrificial layer was patterned and deposited using the same lift-off process. The mechanical support for the biosensors was patterned using negative photoresist (SU-8 2000.5, Kayaku Advanced Materials). Using a thinner (SU-8 2000 thinner, Kayaku Advanced Materials), the SU-8 composition was adjusted to 9.5 % solids to achieve 200 nm SU-8 thickness. The biosensor array interconnects were patterned using similar methods as the outer interconnects. 1nm Cr (99.998%, Kurt J. Lesker Company) / 50nm Pd (99.99%, Kurt J. Lesker Company) / 25 nm Cr (99.998%, Kurt J Lesker) / 10 nm Au (99.999%, Kurt J Lesker) were deposited using an electron beam evaporator (Pro Line PVD Evaporator, Kurt J. Lesker Company). Lastly, the interconnects of the biosensors were passivated with 100 nm of SU-8 2000.5 diluted with SU-8 thinner to 7.15% solids, patterned using photolithography to open the electrode sites (Figure S1).

### 3D-SR-BA structure release

Post fabrication, the 3D-SR-BA chips were cleaned with acetone, IPA, and N2 blow dried. A rectangular chamber was prepared by using 10:1 base:curing agent polydimethylsiloxane (PDMS) (Sylgard 184 Silicone Elastomer, Dow Corning), cured overnight at 65 °C, cut to the needed dimensions and positioned on a device chip, such that the well surrounds the 3D-SR-BA region. A 1% H2O2 (catalog no. 216763, Sigma-Aldrich) solution in deionized (DI) water was added to the well for 1 hour up-to overnight to dissolve the sacrificial layer and trigger self-rolling. After 3D-SR-BAs conformed in 3D, the solution was exchanged for DI water. The radius of curvature was estimated from a slice of 3D confocal microscopy image stack as depicted in Figure S1.

### Poly(3,4-ethylenedioxythiophene): poly(sodium 4-styrenesulfonate) electrodeposition

3D-SR-BA modification with poly(3,4-ethylenedioxythiophene): poly(sodium 4-styrenesulfonate) (PEDOT:PSS) reduces the MEA’s impedance, which results in higher signal amplitudes of ephys recordings.^18^ Electrodeposition was performed in a three-electrode cell setup using a potentiostat (Gamry R600+, Gamry Instruments). A Pt wire, Ag/AgCl electrode and Au microelectrode were used as counter, reference, and working electrodes, respectively. A solution of 0.01 M ethylenedioxythiophene (EDOT)^18^ (97%, catalog no. 483028, Sigma-Aldrich) and 0.02 M poly(sodium 4-styrenesulfonate) (PSS) (catalog no. 243051, Sigma-Aldrich) was prepared in DI water. PEDOT:PSS electrodeposition was performed by using a constant current density of 0.5 mA/cm^2^ applied for 10 min to each electrode sequentially.

### Electrochemical characterization of microelectrodes

To characterize the impedance of the electrodes before and after PEDOT:PSS electrodeposition, electrochemical impedance spectroscopy (EIS) was performed in a three-electrode electrochemical cell using a potentiostat (Gamry R600+, Gamry Instruments).^17^ 1X Phosphate Buffered Saline (PBS) (catalog no. 10010023, ThermoFisher) was used as an electrolyte solution. A Pt wire, an Ag/AgCl electrode and an Au microelectrode were used as counter, reference, and working electrodes, respectively. The frequency was scanned from 10 Hz to 100,00 Hz with VDC of 0 V and VAC of 10 mV. EIS experiments (n = 11) were performed before and after PEDOT:PSS electrodeposition. Measurements were performed in a Faraday cage.

### Cortical neurons isolation

Cortical neurons were obtained from primary embryonic day 18 (E18) rat cortical tissue from BrainBits LLC (SKU: SDECX). For cortical tissue dissociate, 2 mL media from the tissue’s vial was aliquoted and stored for a later step. The tissue was incubated with a solution of 0.25% trypsin ethylenediaminetetraacetic acid (EDTA) (catalog no. 25200056, ThermoFisher) for 5 min. The tissue was again incubated with a solution of 0.25% trypsin EDTA supplemented with 0.1% Deoxyribonuclease (DNase, catalog no. EN0521, ThermoFisher) for another 5 min. Post incubation, the solution was replaced by the previously removed media. The tissue was physically dissociated by triturating it 5 times using a sterile fire-polished glass Pasteur pipette ca. 1 mm in diameter. This was followed by another set of trituration (5 times) with a narrower pipette (ca. 1/2-3/4 mm in diameter). The dissociated cells were then centrifuged at 1100 rpm for 1 min and re-suspended in 1 mL of fresh Neurobasal media (catalog no. A1413701, ThermoFisher) supplemented with 1× B27 (catalog no. A1413701, ThermoFisher), 1% Glutamax (catalog no. 35050-061, ThermoFisher) and 1% Penicillin/Streptomycin (catalog no. 15140122, ThermoFisher).

### Cortical spheroid formation

Cortical spheroids were formed following previously established protocols. ^19–20^ Briefly, sterile 10 g/L agarose solution was prepared by mixing agarose powder (catalog no. 16500, Invitrogen) with a solution of 0.9% (w/v) NaCl (catalog no. S5886, Sigma) in DI water. The mixture was heated using a microwave to completely dissolve the agarose powder. The molten agarose solution was allowed to cool to ca. 60 °C before casting it into an inverse micromold (3D Petri Dish™, Microtissues Inc.) to form spherical microwells as illustrated in Figure S2 A I-II. After solidifying, the 3D Petri Dish with agarose was inverted and agarose microwells were extracted (Figure S2 A III-IV). The agarose microwells were equilibrated twice in Neurobasal medium at 37°C for 30 min in an incubator with 5% CO2. The obtained cortical neurons were seeded in the microwells at a density of 500,000 cells/mold in 190 μL of Neurobasal medium supplemented with 1× B27, 1% Glutamax and 1% Penicillin/Streptomycin (Figure S2 B I-II). The cells settled in microwells by gravity (Figure S2 B III) and started to aggregate. An additional 2 mL of Neurobasal medium was added to the side of each well dropwise (Figure S2 B IV) to avoid disruption of the settled cells. 50% of the cell culture media was replaced with fresh media every 3 days. The cortical neurons self-assembled to form spheroids, which matured in culture for up to 30 days. Four-week-old cortical spheroids were harvested by inverting the micromold (Figure S2 B V). The harvested spheroids were placed in a Tyrode’s solution-filled recording chamber attached to the 3D-SR-BA chip.

### Electrical and optical recording from cortical spheroids

Prior to recording, cortical spheroids were stained with Ca^2+^ indicator, Cal-520 (catalog no. 21130, AAT Bioquest). 5 μM Cal-520 solution was prepared in Tyrode’s solution. Tissues were incubated at 37 °C for 30 min. Post incubation, the tissues were washed 3 times with 37 °C Tyrode’s solution. Each 3D-SR-BA chip was glued to a printed circuit board (PCB) with soldered 36 pin connector (catalog no. A79024-001, Omnetics). The electrodes on the chip were wire-bonded to the PCB using a manual wedge wire bonder (7476D Wire Bonder, West Bond). The chip was mounted on the microscope stage, and a 37 °C temperature inside the measurement chamber was maintained using constant perfusion of heated (Inline heater, ThermoClamp, Automate Scientific) carbogen (5% CO2 in O2 balance, Airgas) bubbled Tyrode’s solution. 50 μM glutamate solution was prepared in Tyrode’s solution, with pH maintained via bubbling of carbogen gas.

For electrical recordings, the Omnetics connector on the printed circuit board was connected to a 32-channel amplifier (RHD2132, Intan Tech.), and the electrical signals were recorded using a commercially available acquisition system (RHD2000, Intan tech.) with a sampling rate of either 20 kHz or 30kHz. Fluorescence imaging was performed using an upright microscope equipped with a resonant confocal scanner (Nikon A1R) with a 20X/0.50 NA water immersion objective. All of the recordings were performed in a grounded Faraday cage.

Four-week old cortical spheroids were transferred to a 3D-SR-BA chip with a temperature - controlled Tyrode’s solution-filled chamber bubbled with carbogen gas. Using an x-y-z micromanipulator (SMX, Sensapex), the arrays were unrolled and the spheroids were encapsulated by a 3D-SR-BA as illustrated by the schematic in Figure 1 A and the confocal microscopy image in Figure 1 C. Functional Ca^2+^ imaging was obtained by confocal microscopy from a single focal plane over a ca. 0.2 mm^2^ field of view, attained at a rate of 59 frames per second.

### Signal Processing and Data Analysis

The recorded neural data from the 3D-SR-BAs was analyzed to eliminate electronic interference and identify single unit neural activity through spike-sorting. Second-order digital infinite impulse response (IIR) notch filters (bandwidth of 5 Hz) were applied with a forward-backward implementation to eliminate 60 Hz power-line interference signal and its harmonics. This postprocessing step was implemented using the open-source SciPy Python library (Python 3.7.2 and SciPy 1.2.1).^21^ Since the spectral content of the recorded neural signal is distributed over 300-7000 Hz frequency range, any localized narrow-band spectral feature can be attributed to unwanted interference, for example, from nearby electronic equipment or the power line. In our analysis, we detected narrow-band interference in the frequency domain by applying a peak-detection algorithm on the signal power spectral density (PSD). Peaks were identified from the peak-prominence, defined as the vertical distance between the peak and its lowest distinguishable contour line in the frequency domain. A peak-prominence threshold of 0.035 was empirically chosen to minimize distortion to the neural data while targeting prominent narrow-band interference. These identified frequency peaks were subsequently removed from the recorded data using the same IIR notch filter settings used to remove 60 Hz harmonics. Additionally, a bandpass filter from 300 Hz to 7 kHz was applied using Python and the open-source SpikeInterface library to limit the spectral component of the signal to the spike band.^22^

Single unit neural activity was extracted from the filtered data using Python (SpikeInterface library) and the MountainSort4 spike-sorting algorithm.^23^ Spike events were first detected using a threshold of 5.0 standard deviations away from the mean of the recorded signal. Dimensionality reduction was performed on the events using principle component analysis (PCA) with n=3 components on each recording channel. Then, the detected events were clustered into separate neural units based on the PCA results using the MountainSort4 algorithm.^24^ The neural unit clusters were manually curated by inspecting the average spike waveforms and inter-spike intervals for each unit cluster to remove noise and recording artifacts misidentified as neural units. Firing rate histograms were calculated from the spike event time information using a bin size of 2 seconds.

## Results and Discussion

### Fabrication and Electrochemical Characterization of Biosensors on 3D-SR-BA

3D-SR-BA is realized using a pre-stressed metal/polymer multilayer structure as previously reported.^15^ Our self-rolling platform is fabricated on a planar surface (Figure 1 A I, II) on top of a sacrificial layer and can accommodate multiple sensor types.^15^ Upon dissolution of the Ge sacrificial layer in 1% hydrogen peroxide, the pre-stress in the metallic traces guides the array’s self-rolling, achieving a controlled 3D sensor configuration with a desired curvature (Figure 1 A I, II, and Figure S3, Materials and Methods).^15^ The curvature of the 3D-SR-BAs is controlled by modulating the polymer insulation layer thickness, resulting in highly-tunable arrays with controllable radius of curvature.^15^ In this study, we fabricated the 3D-SR-BA (radius of curvature 96±4 μm, Figure 1 B, Figure S3) to match the dimensions of a cortical spheroid (diameter 202 ±24 μm, Figure 1 C, Figure S4).

Microfabricated biosensors, such as MEAs, are used extensively in neural interfaces^25^ due to their potential for long-term, multisite, and simultaneous recordings with high temporal resolution.^26^ Microfabrication techniques allow tailoring the array geometry, size and substrate mechanical properties (e.g. fabrication of ultra-compliant probes^25, 27–31^). In this study, seventeen 25 μm ˟ 25 μm microelectrodes are fabricated on the 235 μm ϗ 530 μm 3D-SR-BA (Materials and Methods) and assembled in 3D to interrogate the electrical activity of cortical spheroids with cellular resolution, as illustrated by the 3D confocal image (Figure 1 B). To reduce impedance and produce higher signal amplitudes during recording,^18^ the Au microelectrodes are coated with with PEDOT:PSS via electrodeposition post fabrication.^18^ PEDOT:PSS modification resulted in a reduction in microelectrode impedance from 1.43 ± 0.40 MΩ prior to modification (Figure S5, blue trace, measured at 1 kHz, n = 11) to 21.3 ± 3.5 kΩ after electropolymerization (Figure S5, orange trace, measured at 1 kHz, n = 11).^15, 18^ The reduced impedance results from improved capacitance due to the electrical double layer formation at the interconnected PEDOT-rich and PSS-rich grains.^32^

### Electrical and Optical Recording of Spontaneous Firing of Neurons in Cortical Spheroids

3D electrophysiological signals were recorded from cortical spheroids using 3D-SR-BAs. Cortical spheroids were compacted in agarose microwells for 28 days. Rod-shaped cortical tissues were transferred to the 3D-SR-BA chip with a temperature-controlled carbogenated Tyrode’s solution-filled chamber (Materials and Methods). Using an x-y-z micromanipulator, the 3D-SR-BAs were unrolled allowing each array to encapsulate a single spheroid as illustrated in Figure 2 A. Six microelectrodes arranged in 3D (Figure 2 B) simultaneously recorded the field potentials from the encapsulated cortical spheroid (Spheroid 1). The 3D-SR-BA electrical recordings were obtained concurrently with functional Ca^2+^ imaging (Figure 2 C, Supplementary Movie 1). Ca^2+^ ions flow across the cell membrane during an action potential, allowing optical monitoring of the resulting electrical activity via fluorescence imaging.^33^ Functional Ca^2+^ transients (Figure 2 A, green fluorescence, Figure 2 C, Supplementary Movie 1) were simultaneously imaged from a single focal plane over a ca. 0.2 mm^2^ field of view. The spheroid was stained with Ca^2+^ indicator (Cal-520, Materials and Methods). Cortical neurons cultured *in vitro* for an extended time exhibit synchronized neural activity.^34^ This result was confirmed by comparing the functional Ca^2+^ transients between individual region of interest (ROI) and the whole field of view (Figure S6). The Ca^2+^ transients acquired from the whole field of view captures all the transients from the ROIs.

**Figure 2.**
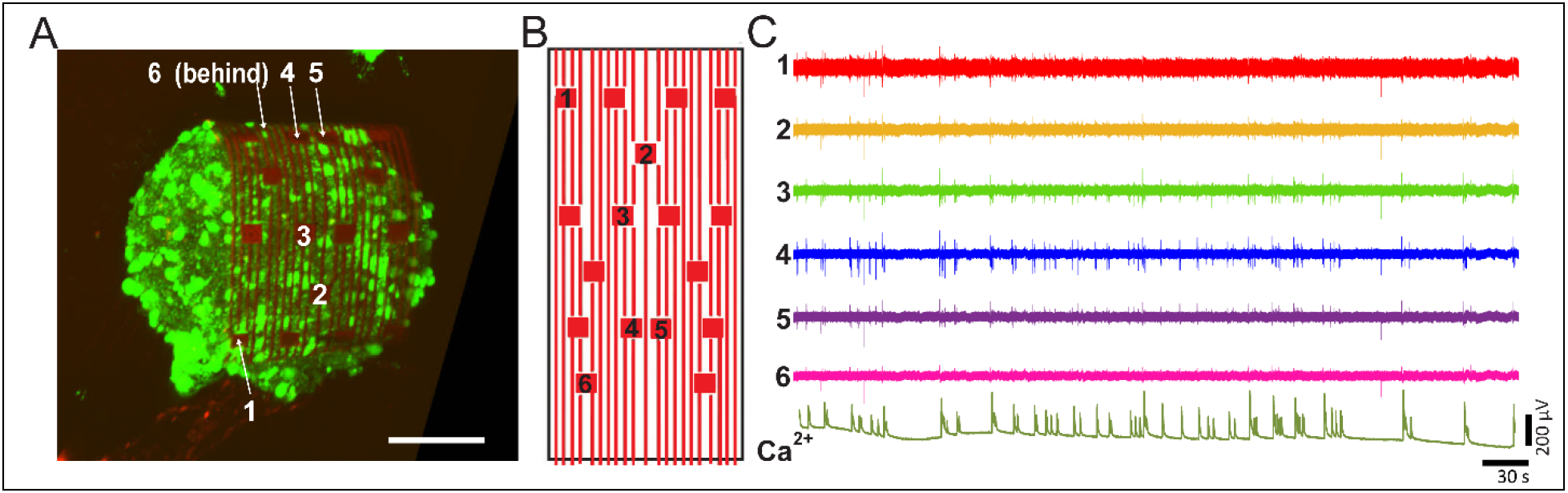
Electrical and optical recordings in 3D of cortical spheroids using 3D self-rolled biosensor array (3D-SR-BA). **(A)** A 3D confocal microscopy image of 3D cortical spheroid labeled with Ca^2+^ indicator dye (Cal-520, green fluorescence) encapsulated by the 3D-SR-BA. Scale bar is 100 μm. **(B)** A 2D map of the recording microelectrodes labeled in panel (A). **(C)** Representative voltage traces recorded from spontaneously firing cortical neurons via channels labeled in panel (A) and (B) with simultaneously recorded Ca^2+^ fluorescence intensity (whole tissue).

The raw neural data recorded from six electrodes was analyzed (Materials and Methods) and five distinct single unit spikes were identified in cortical Spheroid 1. The averaged waveforms of all sorted spikes associated with each unit are illustrated in Figure 3 A (solid color traces). The standard deviations of individual spike events in each unit are represented by the shaded regions in Figure 3 A. Different unit waveforms are detected by different microelectrodes near the firing neurons in the neural spheroid. The corresponding electrode is identified by a number on the left of each waveform plot in Figure 3 A. Three out of the five units were primarily detected on individual electrode channels: each unit on individual channel, indicating that the responsible neurons were close to each of those electrodes. Units 3 and 4 were detected on both channels 4 and 5 (Figure 3 A), indicating that the firing neurons were located between those electrodes (Figure 2 A, B).

**Figure 3.**
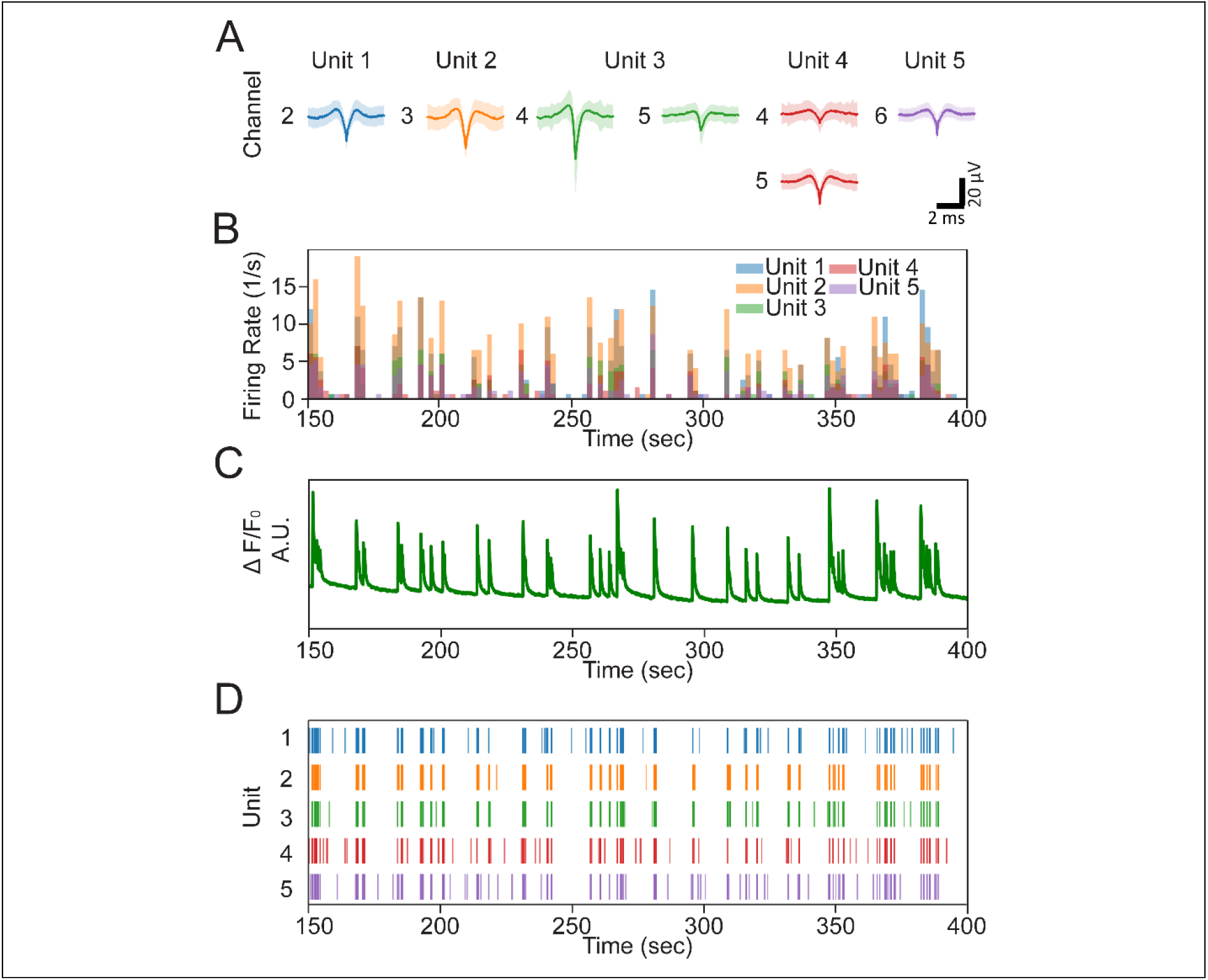
Spontaneous cortical neuron activity correlated to Ca^2+^ fluorescence transients. **(A)** Averaged spike (solid trace) and ± standard deviation (shaded region) detected in each unit. The number on the left of the trace is the channel where the unit was detected. **(B)** Overlaid firing rate of representative units. **(C)** Simultaneously recorded Ca^2+^ fluorescence intensity. **(D)** Raster plots of the firing patterns of the detected units.

The firing rates of individual spike-sorted neural units (Figure 3 B and Figure S7 A) indicated bursts of activity up to 20 spikes/s, which corresponds well with the fluorescent signal of the functional Ca^2+^ imaging (Figure 3 C, Figure S7 B, Supplementary Movie 1). The precise spike timings are shown in the raster plot in Figure 3 D. It is evident that the firing patterns of the detected units are synchronized across the spheroid, which agrees with previously published studies on synchronization of cortical neuron clusters cultured *in vitro*.^34^ This synchrony is also observed in the Ca^2+^ transients obtained from different regions of the spheroid (Figure S6).

### Electrical and Optical Recording of glutamate-induced firing of neurons in cortical spheroids

As a proof-of-concept for 3D-SR-BA use in drug effect monitoring, the effect of glutamate, the main excitatory neurotransmitter of the nervous system,^35^ was investigated. The effect of glutamate on glutamatergic neurons is well established.^35^ The 3D-SR-BA was used to continuously record ephys activity while glutamate solution (50 μM) was perfused into a chamber containing the encapsulated spheroid. Simultaneous functional Ca^2+^ imaging was used to supplement the ephys recording and illustrated the same cellular activity (Figure 4 A, Supplementary Movie 2, Materials and Methods). The same five neuronal units isolated during spontaneous neural activity recording (Figure 4 B) were detected by the data analysis pipeline. Both the electrical and optical recordings show elevated neuronal firing rates following the introduction of glutamate (Figure 4 A, C, D, Figure S8, and Supplementary Movie 2). The increased firing rate is the expected after the addition of excitatory neurotransmitter glutamate.^35^ We note that the time delay between the onset and offset of glutamate flow and the change in neuronal firing rate is due to the time required for glutamate to perfuse into the recording chamber. The raster plots of the detected units illustrate a firing rate increase between 4.5-fold and 9.7-fold for individual units (above the spontaneous activity baseline level) upon addition of glutamate (Figure 4 E, Figure S8).

**Figure 4.**
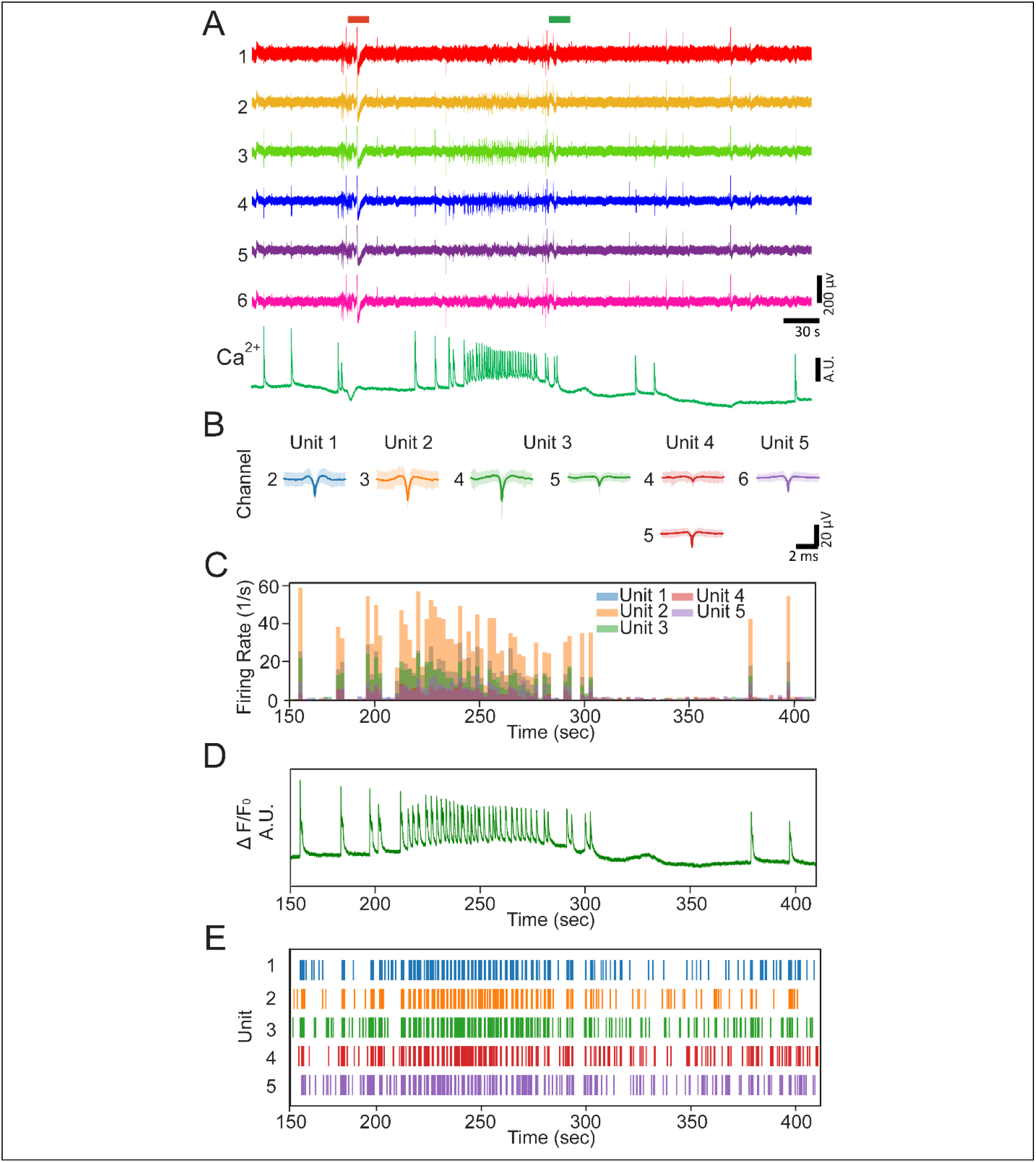
Glutamate-induced activity of cortical neuron is correlated to Ca^2+^ fluorescence transients. **(A)** Representative ephys recording of glutamate-induced cortical neuron activity and corresponding Ca^2+^ fluorescence transients. **(B)** Averaged spike (solid trace) and ± standard deviation (shaded region) detected in each unit. The number on the left of the trace is the channel where the unit was detected. **(C)** Overlaid firing rate of representative units. **(D)** Simultaneously recorded Ca^2+^ fluorescence intensity. **(E)** Raster plots of the firing patterns of the detected units.

The 3D recording demonstrated in this work was repeated by measuring the spontaneous and glutamate-induced firing from three independent neural spheroids (Technical Note 1 in Supplementary Information). The change in spike counts upon glutamate addition are summarized for each spheroid in Figure S9. A notable increase in spiking activity of individual units within a spheroid upon glutamate addition was observed (4.5-9.7-fold, 6.1-14.5-fold, and 2.5-37.9-fold for units in Spheroid 1, 2 and 3, respectively). This increase is associated with the glutamate flow indicated by the green-shaded regions in Figure S9. For each spheroid, we introduced glutamate in multiple cycles and after each cycle glutamate was washed from the chamber. The change in neuronal firing rate (increase upon addition and decrease upon removal) consistently followed the glutamate cycles. The variation of neuronal response to glutamate among different units in each spheroid, characterized as the change in firing rates, can be attributed to biological variability in the spontaneous activity of each unit: neurons that fire more sparsely under the spontaneous condition illustrate a more pronounced increase under the glutamate-perfused condition.

#### Conclusion and future directions

This work demonstrates the first 3D ephys recording from a neuronal spheroid using a 3D biosensor platform. The novel architecture of the 3D-SR-BA allows for high-resolution recordings of neural activity currently unattainable by other recording methods. It provides a robust, tissue incorporated, continuous, sensitive, and reproducible electrical recordings. Integrating a signal-processing and spike sorting analysis pipeline enables isolation of individual distinct neural units from the 3D-SR-BA recorded data. The electrical recordings were supplemented with functional Ca^2+^ imaging and firing rates of the individual units detected with the 3D-SR-BA corresponded to the Ca^2+^ transients imaged optically. Using our 3D-SR-BA neural spheroid system, we are able to measure the spheroid’s response to glutamate addition which results in increased neuronal activity as indicated by the difference in the firing pattern compared to the spontaneous activity recording. The ability to record complex neural electrical activity using 3D-SR-BA opens a plethora of possibilities for longer-term study of 3D neuronal tissues. The approach presented in this work can enable studies of neuronal functions in 3D, e.g. investigation of brain organoid development and functionality in both healthy and disease models.^12^ Moreover, our approach can assist with the discovery of new therapeutics for neurological disorders using an *in vitro* model which is a better approximation of *in vivo* conditions.

## Acknowledgments

T.C.-K. acknowledges funding support from the National Science Foundation (NSF) under Award CBET1552833, the Office of Naval Research under Award N000141712368, and the Defense Advanced Research Projects Agency under Award AWD00001593 (416052-5). M.C. acknowledges funding support from the NSF under Award NCS-FO1926804. A.K. acknowledges support from the Neil and Jo Bushnell Fellowship in Engineering. We also acknowledge support from the Department of Materials Science and Engineering Materials Characterization Facility supported by Grant MCF-677785. J.W.R. acknowledges support by the Carnegie Mellon University Ben Cook Presidential Graduate Fellowship, the Carnegie Mellon University Richard King Mellon Foundation Presidential Fellowship in the Life Sciences, the Axel Berny Presidential Graduate Fellowship, and Philip and Marsha Dowd.

## Supplementary Information

### Technical Note 1: Experimental Replication in Additional Cortical Spheroids

The 3D ephys recording and functional Ca^2+^ imaging demonstrated in this work was repeated under spontaneous and glutamate-perfused conditions for three separately prepared cortical spheroids.

### Spheroid 2

A second spheroid stained with a Ca^2+^ indicator (green fluorescence) was encapsulated by the 3D-SR-BA (Figure S10 A). The recording channel locations are identified by the labels in Figure S10 A, B. The recorded voltage traces are shown alongside the Ca^2+^ fluorescence intensity versus time for the spontaneous (Figure S10 C, Supplementary Movie 3) and glutamate-perfused (Figure S10 D, Supplementary Movie 4) conditions. This recording had several pronounced artifacts, potentially emerging from the initiation of the glutamate flow or other environmental noise (marked by asterisks on Figures S10 C, D). The addition of glutamate (green bars, Figure S10 D) produces a notable increase in ephys and Ca^2+^ activity. This cortical spheroid was subjected to three glutamate addition/wash-out cycles, demonstrating a repeatable glutamate response.

Spike-sorting of the recorded spontaneous ephys activity from this spheroid revealed three distinctly clustered units (Figure S11 A). The firing rates of these units illustrated on Figure S11 B and Figure S12 A, are supplemented by the simultaneously recorded functional Ca^2+^ imaging (Figure S11 C, Figure S12 B, Supplementary Movie 3). The spontaneous firing is sparse for this neuronal construct, shown in the raster plots for individual units (Figure S11 D). Upon glutamate addition, the number of active detected units in the second spheroid increases to eight (Figure S13 A). This potentially indicates glutamate stimulation of glutamatergic cortical neurons that are otherwise dormant.^36^ Several units were identified across multiple channels, such as units 5 and 7, suggesting that they are located between the two electrodes. The glutamate addition resulted in increased spiking activity, as evidenced by the increased firing rates (Figure S13 B, Figure S14 A, and Supplementary Movie 4), which reverted upon glutamate removal. The onset of Ca^2+^ transients aligns with the firing of individual units (Figure S13 B, C, D and Figure S14). The introduction of glutamate induced a 6.1-fold increase to 14.5-fold increase in the firing rate of individual neurons.

### Spheroid 3

Using the same method, the third spheroid was labeled with a Ca^2+^ indicator and encapsulated by the 3D-SR-BA (Figure S15 A). The electrical activity of this spheroid was recorded by the channels marked in Figure S15 A, B. The ephys recordings and functional Ca^2+^ imaging were obtained for the spontaneous (Figure S15 C, Supplementary Movie 5) and the glutamate-induced neuronal activity (Figure S15 D, Supplementary Movie 6). An artifact, potentially from the initiation of the glutamate flow, is marked by an asterisk on Figure S15 D. The neural activity measured by ephys recordings and functional Ca^2+^ imaging increased after glutamate addition and decreased upon glutamate removal. This spheroid was subjected to two glutamate addition and removal cycles, demonstrating the expected glutamate response in each cycle.

Six distinctly clustered units were identified in the spontaneous recording (Figure S16 A). The firing rates of these units illustrated in Figure S16 B and Figure S17 A, are supplemented by the simultaneously recorded functional Ca^2+^ imaging (Figure S16 C, D, Figure S17 B, and Supplementary Movie 5). Units 3 and 4 were identified across multiple channels, suggesting that they are positioned between those electrodes. Upon glutamate addition, however, the number of detected units in spheroid 3 decreases to four (Figure S18 A). Several units were still identified across multiple channels, such as units 2 and 4, suggesting that the responsible neuron is located between the electrodes. The glutamate addition resulted in increased spiking activity, as evidenced by the increased firing rates (Figure S18 B, Figure S19 A, and Supplementary Movie 6), which reverted upon glutamate removal. The onset of Ca^2+^ transients aligns with increased firing of individual units (Figure S18 B, C, D and Figure S19). The introduction of glutamate induced an increase between 2.5-fold and 37.9-fold in firing rate of individual neurons within the spheroid.

### Results Across Multiple Spheroids

The data indicate that in some cases (as in Spheroid 2), the number of detected units is higher after the introduction of glutamate, whereas in others (as in Spheroid 3), fewer units are detected in the glutamate condition. The addition of glutamate could excite different subsets of glutamatergic neurons which only partially overlaps with the population of spontaneously active neurons. Alternatively, the detection of specific units could also be an artifact of the spike-sorting, whereby the varied distribution of neural activity under glutamate stimulation lends itself to better or poorer single-unit isolation for certain neurons.

The change in spike counts are summarized for each condition in Figure S9, illustrating a notable increase in spike count from spontaneous to glutamate-induced firing in each of the recordings. Such an increase is evidently associated with the glutamate flow specified by the green-shaded regions in Figure S9. The notable increase of neuronal firing upon multiple cycles of glutamate addition and decrease in firing upon glutamate removal indicates a predicted and repeatable response in this biological system.

---

**Figure S1.**
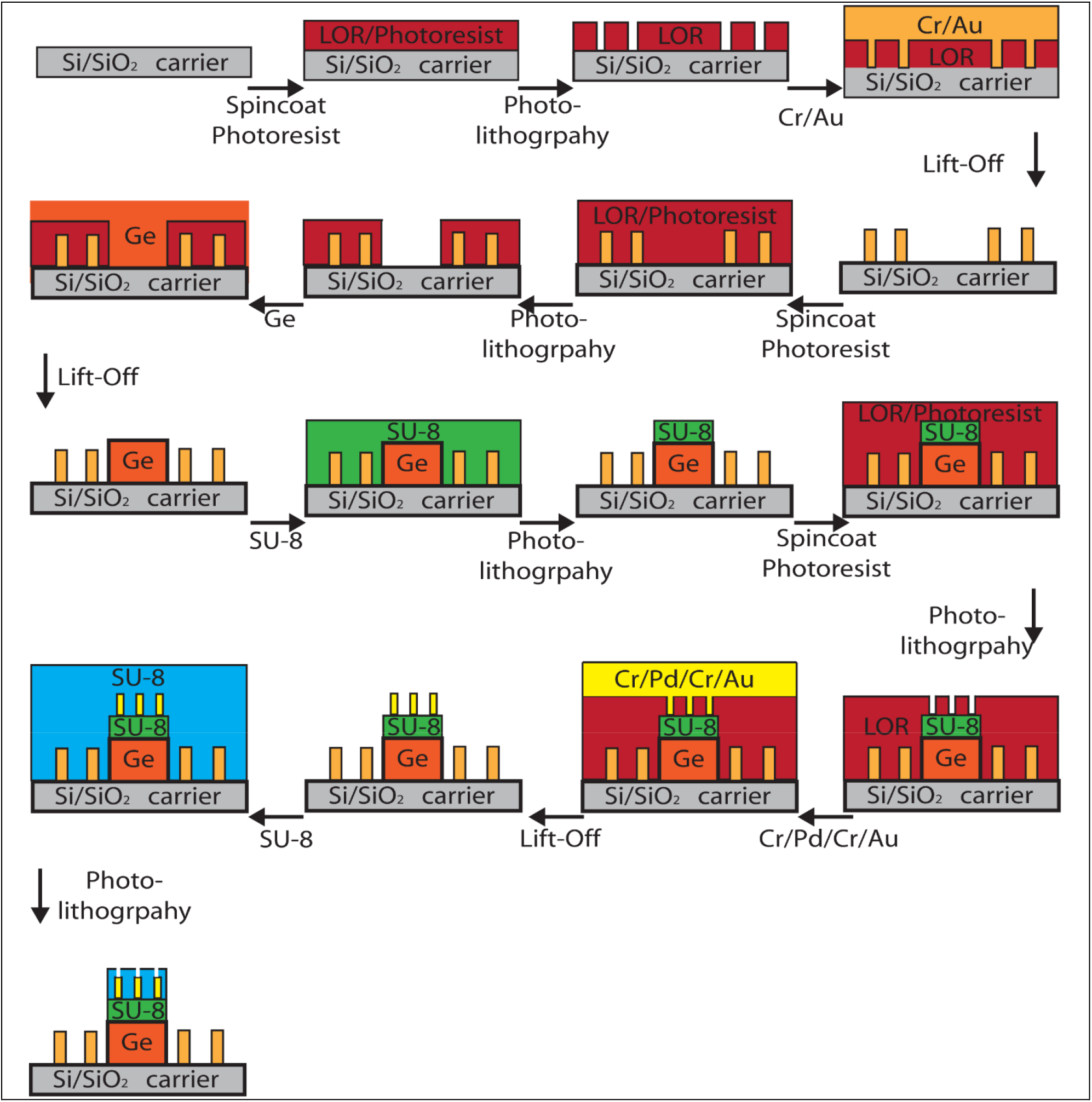
Fabrication of 3D-SR-BAs. Schematics representing the fabrication flow process. The following are the main steps: outer electrodes interconnects (Cr/Au), Ge sacrificial layer, SU-8 base layer, biosensor and interconnects (Cr/Pd/Cr/Au), followed by SU-8 passivation layer.

**Figure S2.**
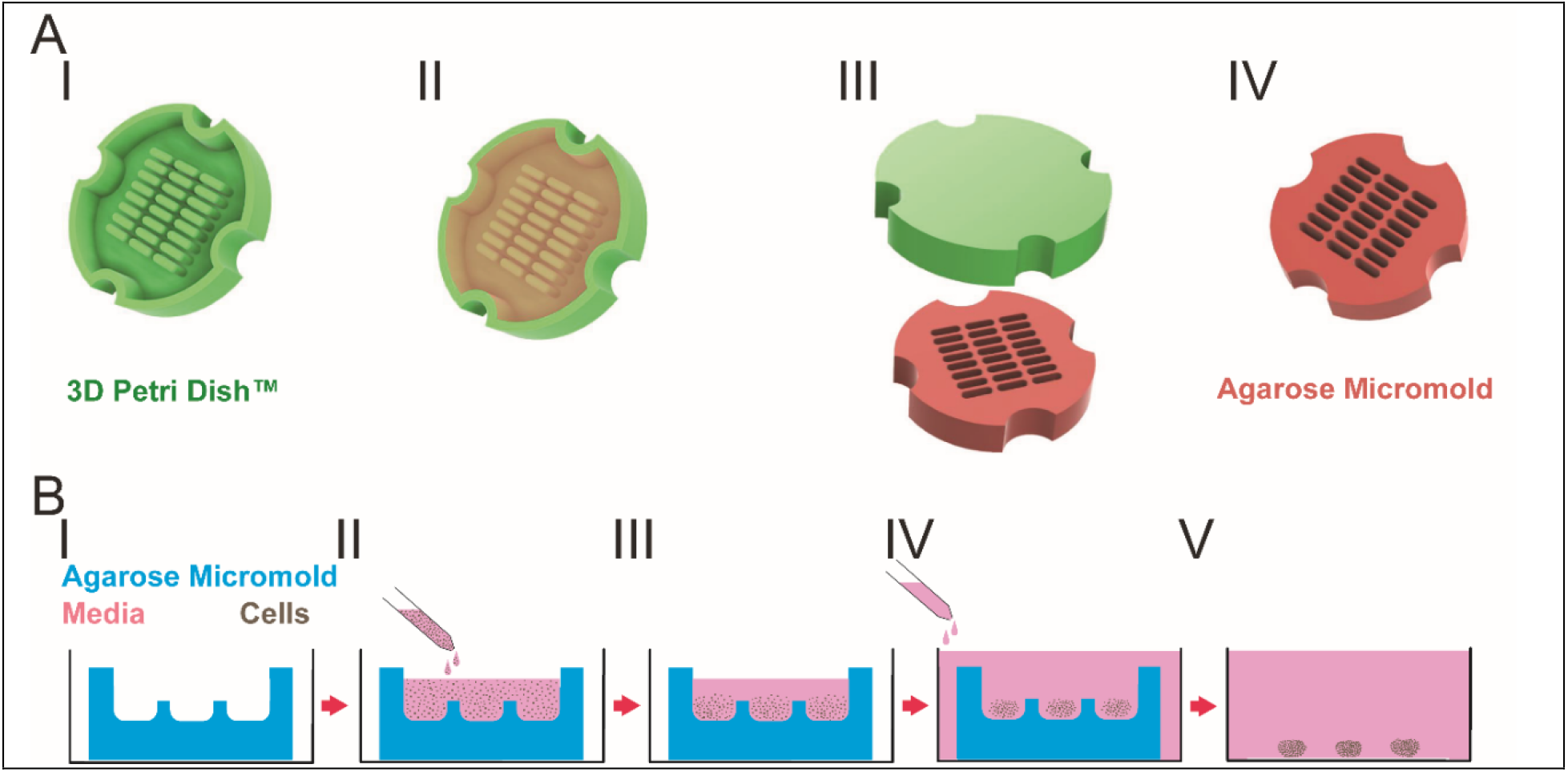
Spheroid formation using 3D Petri Dish Micromold Microtissue™. **(A)** A schematic of casting an agarose microwell using 3D Petri Dish™ (**I**) Sterilized 3D petri dish is (**II**) filled with molten agarose, bubbles are removed, and complete gelation of agarose is achieved. (**III**) The 3D Petri Dish™ is inverted and flexed to remove the cast agarose microwells. (**IV**) Agarose micromolds are equilibrated in media prior to seeding. **(B)** A schematic for cell seeding and spheroid formation. (**I**) The equilibrated agarose micromold is placed in a well/plate, excess medium is removed. (**II**) Cell suspension in media is seeded on top of the micromold dropwise. (**III**) Cells settle into microwells by gravity. (**IV**) Culture media is added to the well/plate outside the agarose micromold. Cells compact and form spheroids over time. (**V**) The compacted spheroids harvested from the agarose micromold.

**Figure S3.**
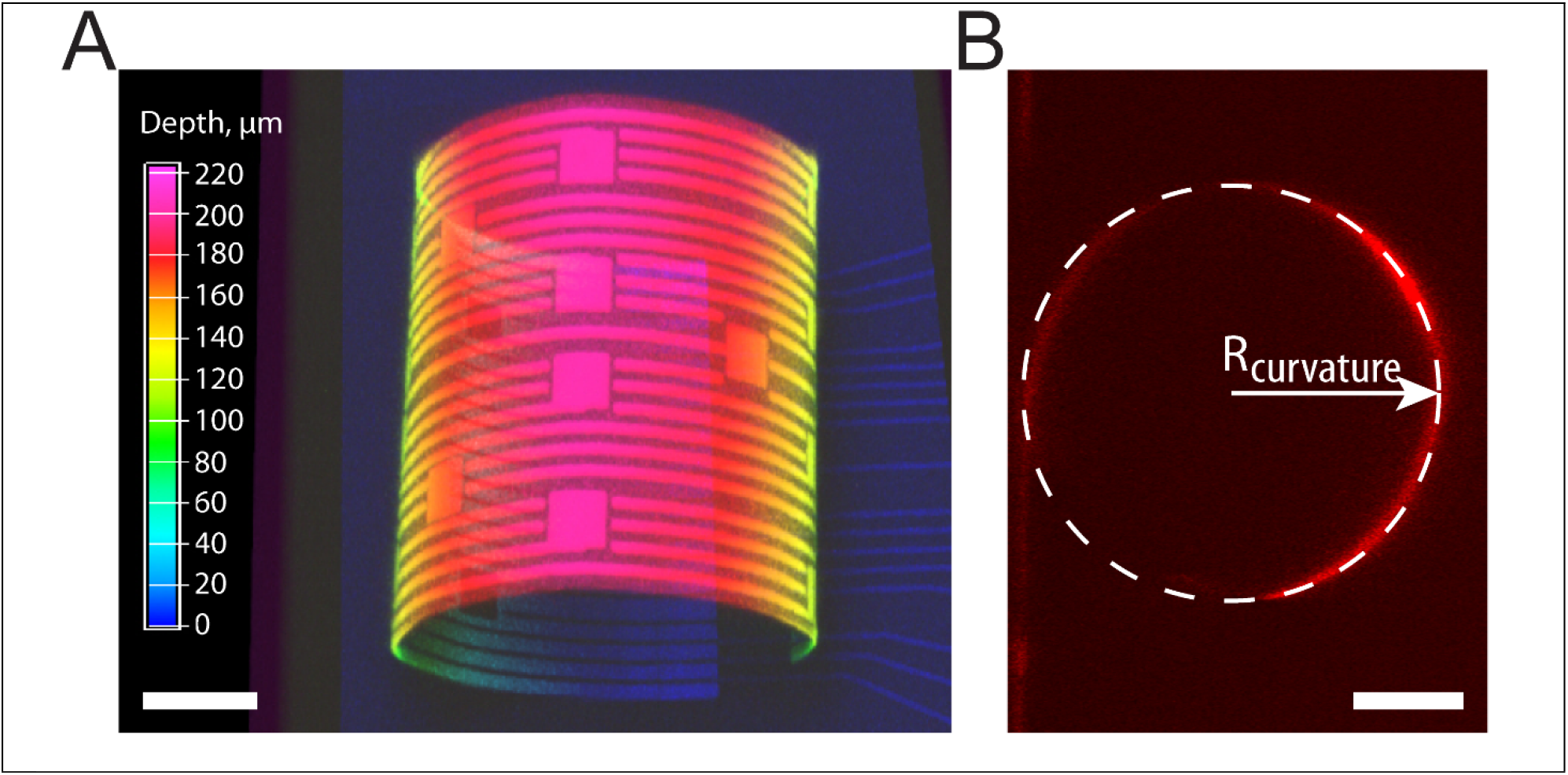
Estimation of the radius of curvature for 3D self-rolled biosensor array (3D-SR-BA). **(A)** A 3D confocal microscopy image of fabricated 3D-SR-BA with ~1 turn. Color bar represents the depth in μm, scale bar is 50 μm. **(B)** Measurement of the radius of curvature (dashed while circle) from a 3D confocal microscopy image slice of 3D-SR-BAs with ~1 turn, scale bar is 50 μm.

**Figure S4.**
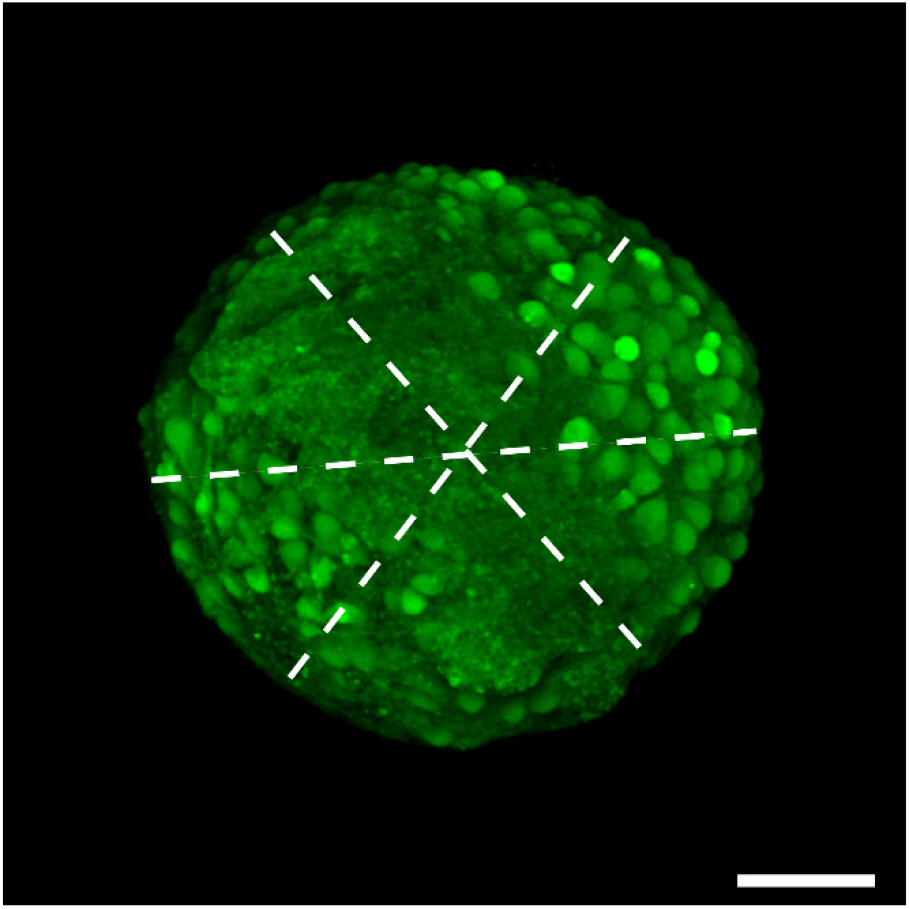
Estimation of the diameter of a rat cortical spheroid. Diameter measurements (dashed white lines) of a representative cortical spheroid stained with calcein (green fluorescence) obtained from a maximum intensity projection of 3D confocal microscopy images. Scale bar is 50 μm.

**Figure S5.**
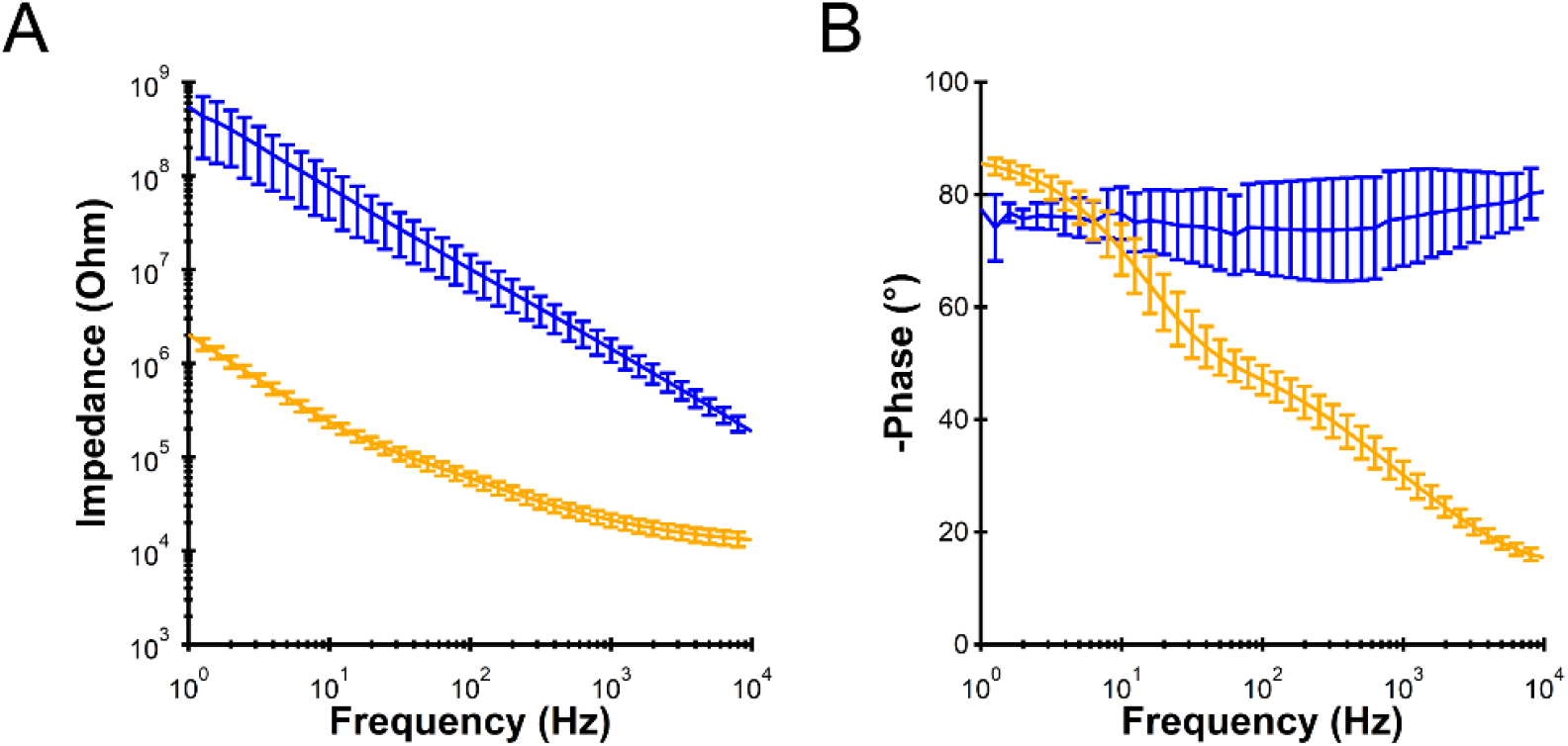
Electrochemical impedance spectroscopy (EIS) characterization of the electrodes before (blue trace) and after (orange trace) PEDOT: PSS electrodeposition. **(A)** Magnitude of the impedance and **(B)** Negative phase as a function of frequency. Data is presented as mean ± SD (n = 11 electrodes).

**Figure S6.**
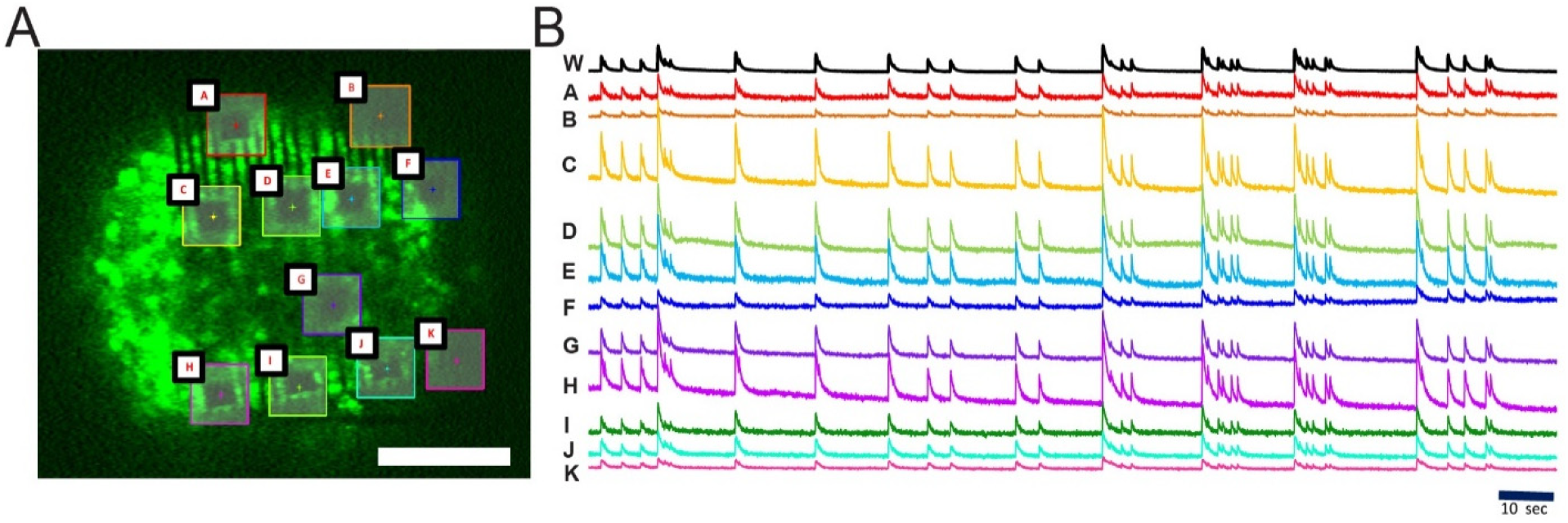
Functional Ca^2+^ imaging from local regions of interest on cortical spheroid correspond to the whole-tissue optical recording of Ca^2+^transients. **(A)** A 3D confocal microscopy image of 3D cortical spheroid labeled with Ca^2+^ indicator dye (Cal-520, green fluorescence) encapsulated by the 3D self-rolled biosensor array (3D-SR-BA). **(B)** Representative Ca^2+^ transients from the regions of interest marked by boxes in panel (A). “W” denotes whole-tissue fluorescence. Scale bars are 100 μm.

**Figure S7.**
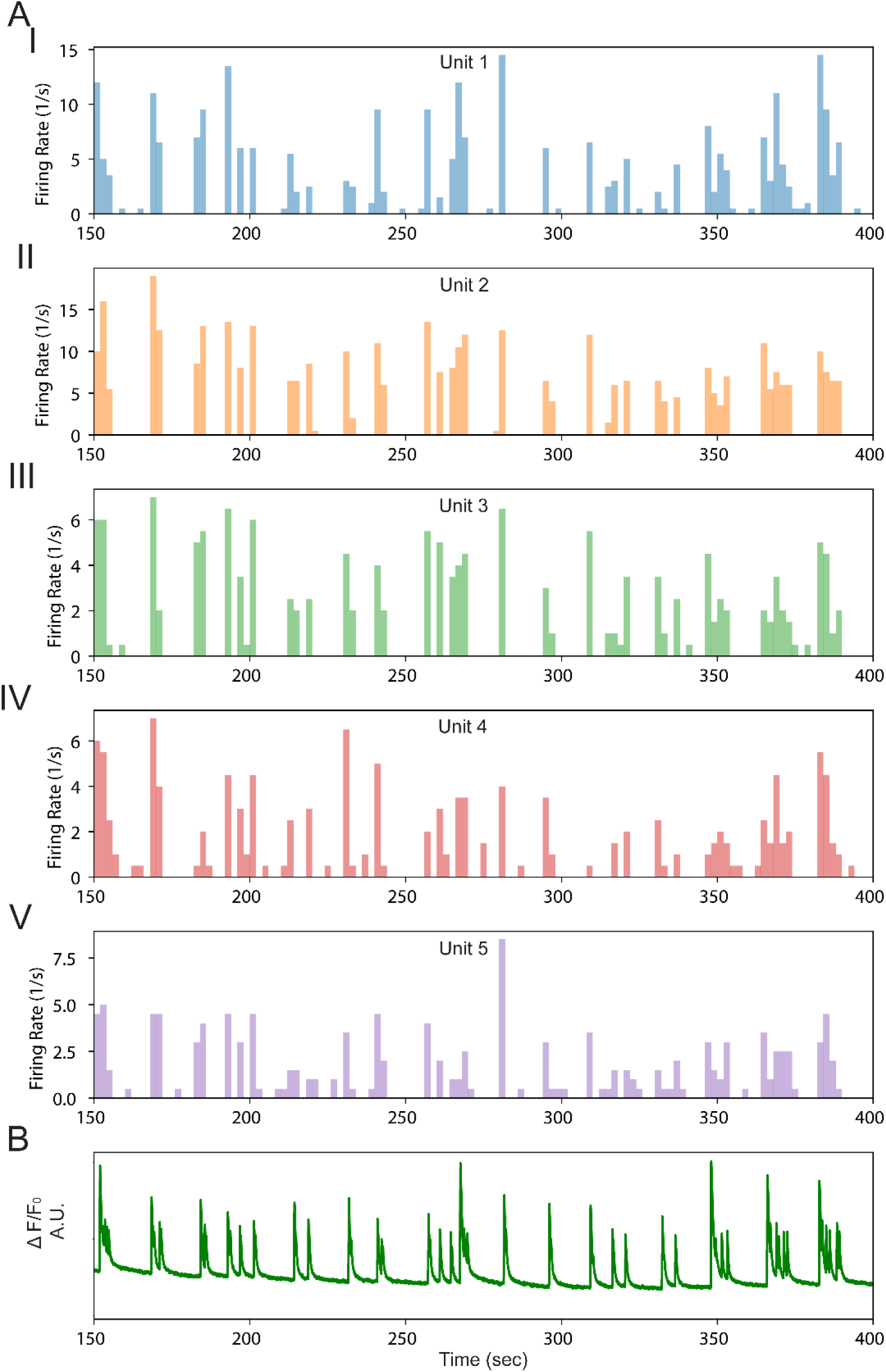
Spontaneous firing of representative units detected after spike sorting of the field potential recording and corresponding whole-spheroid Ca^2+^ fluorescence. **(A)** Firing rate of individual units: (**I**) unit 1, (**II**) unit 2, (**III**) unit 3, (**IV**) unit 4, and (**V**) unit 5. **(B)** Optically recorded Ca^2+^ fluorescence intensity from the same time range.

**Figure S8.**
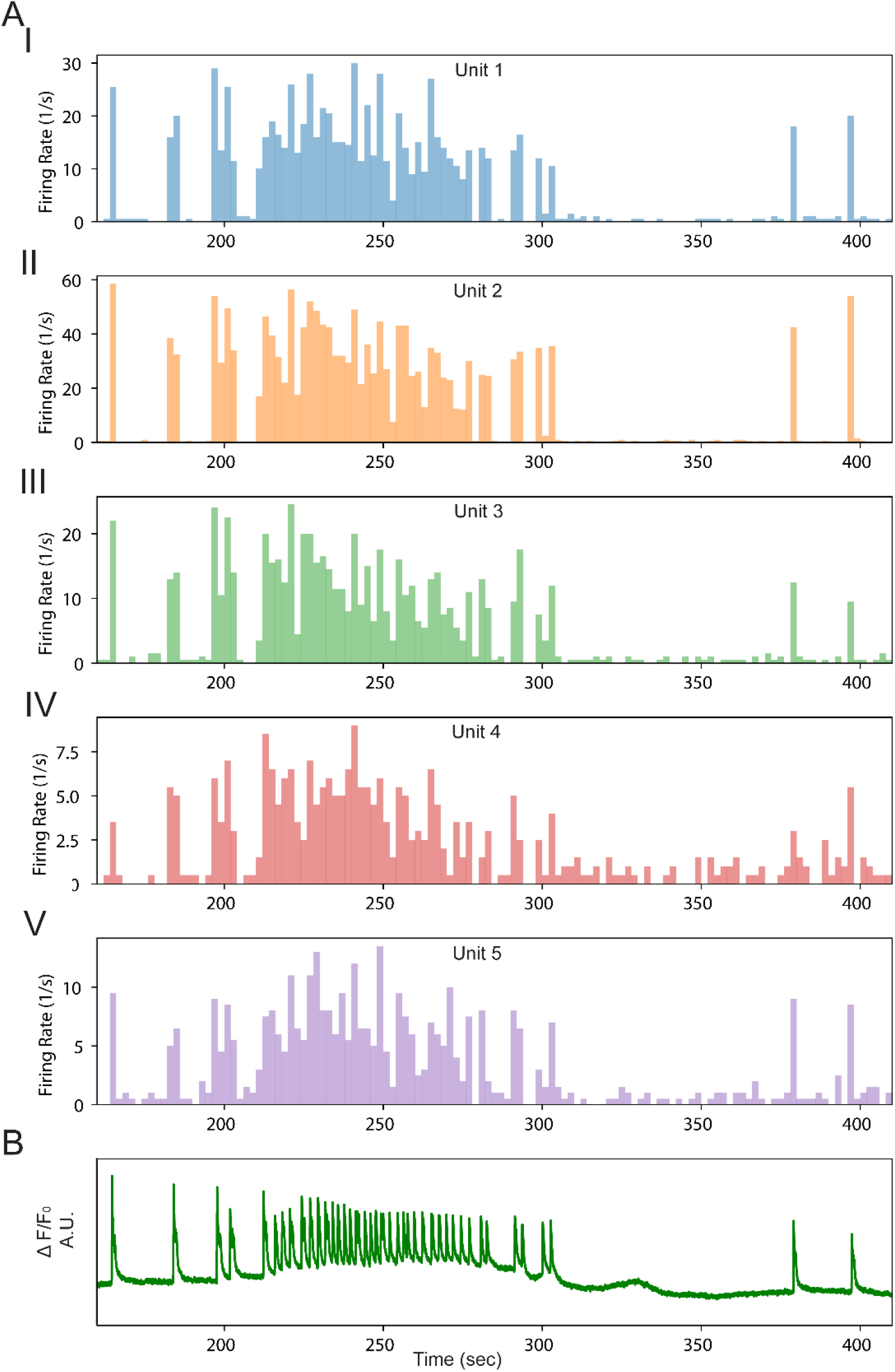
Glutamate-induced firing of units detected after spike sorting of the field potential recording and corresponding whole-spheroid Ca^2+^ fluorescence. **(A)** Firing rate of individual units: (**I**) unit 1, (**II**) unit 2, (**III**) unit 3, (**IV**) unit 4, and (**V**) unit 5. **(B)** Optically recorded Ca^2+^ fluorescence intensity from the same time range.

**Figure S9.**
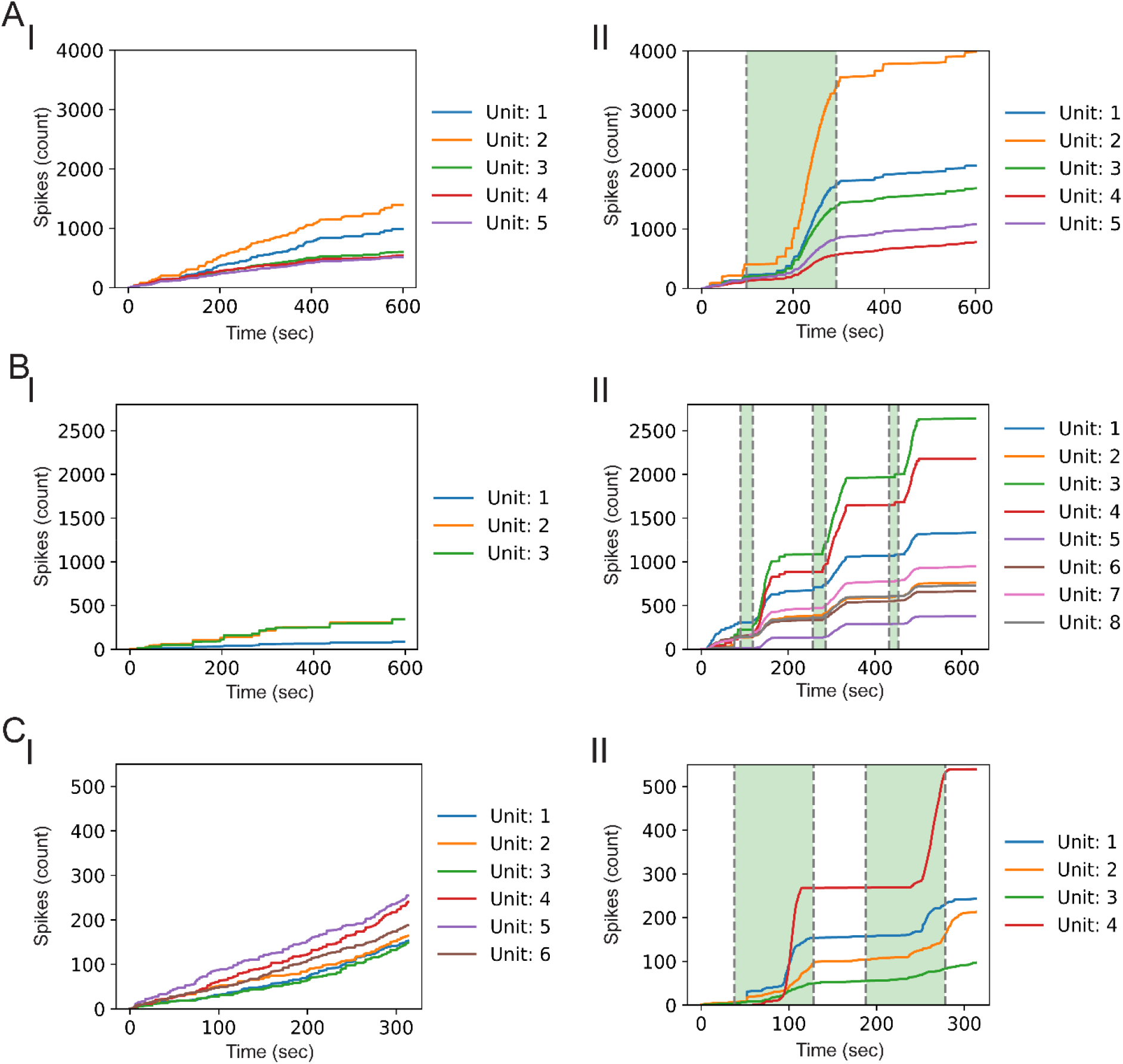
Cumulative sum plot of firing rates of representative units detected after spike sorting of the field potential recording. **(A) Spheroid 1**(**I**) spontaneous activity, (**II**) glutamate-induced activity. **(B) Spheroid 2**(**I**) spontaneous activity, (**II**) glutamate-induced activity. **(C) Spheroid 3**(**I**) spontaneous activity, (**II**) glutamate-induced activity. Shaded green region indicates glutamate flow.

**Figure S10.**
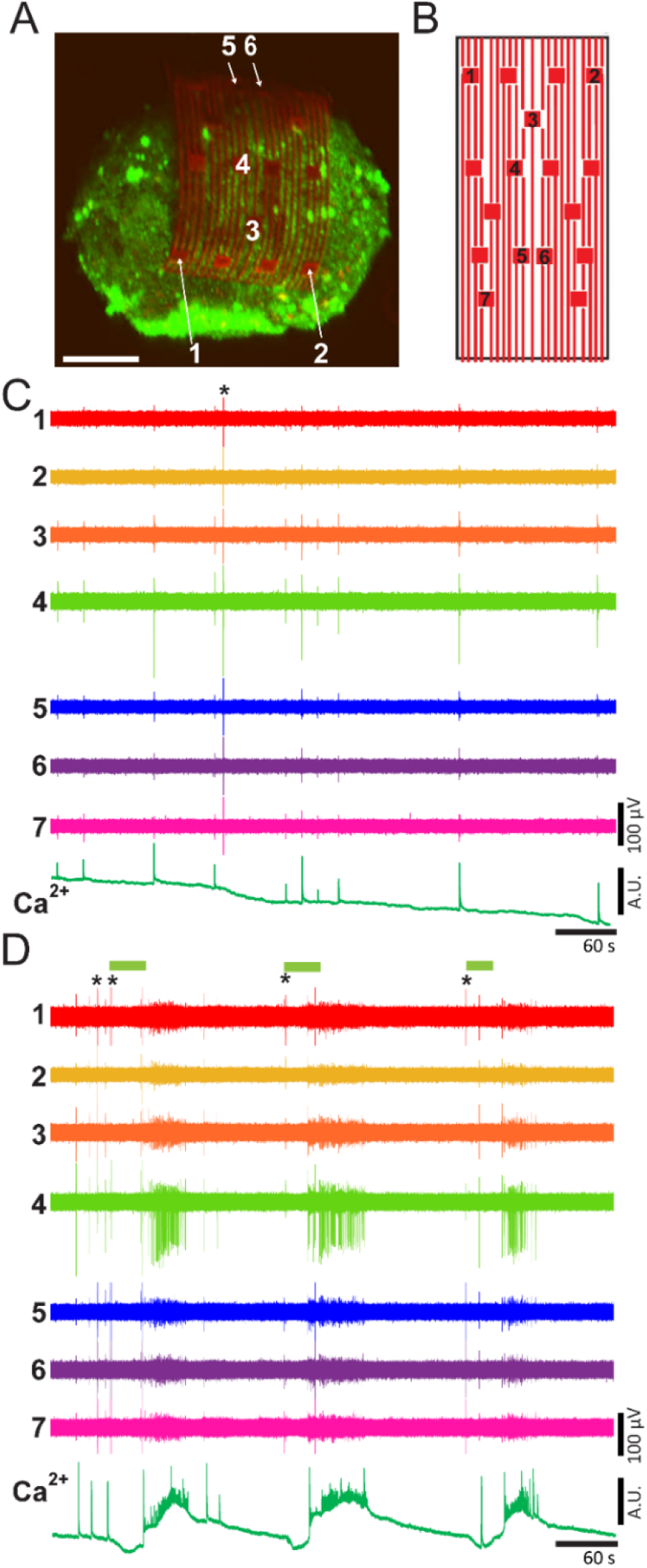
Electrical and optical recordings in 3D of cortical spheroids (second spheroid) using 3D-self-rolled biosensor array (3D-SR-BA). **(A)** A 3D confocal microscopy image of 3D cortical spheroid labeled with Ca^2+^ indicator dye (Cal-520, green fluorescence) encapsulated by the 3D-SR-BA. Scale bar is 100 μm. **(B)** A 2D map of the microelectrodes labeled in panel (A) recording electrical activity of cortical neurons. **(C)** Voltage traces of spontaneously firing cortical neurons recorded from the channels labeled in panel (A) and (B) with simultaneously recorded Ca^2+^ fluorescence intensity (whole tissue). **(D)** Voltage traces of cortical neurons firing upon addition of glutamate recorded from the channels labeled in panel (A) and (B) with simultaneously recorded Ca^2+^ fluorescence intensity (whole spheroid). Green bars indicate glutamate flow. **\*** Denotes electrical artifacts.

**Figure S11.**
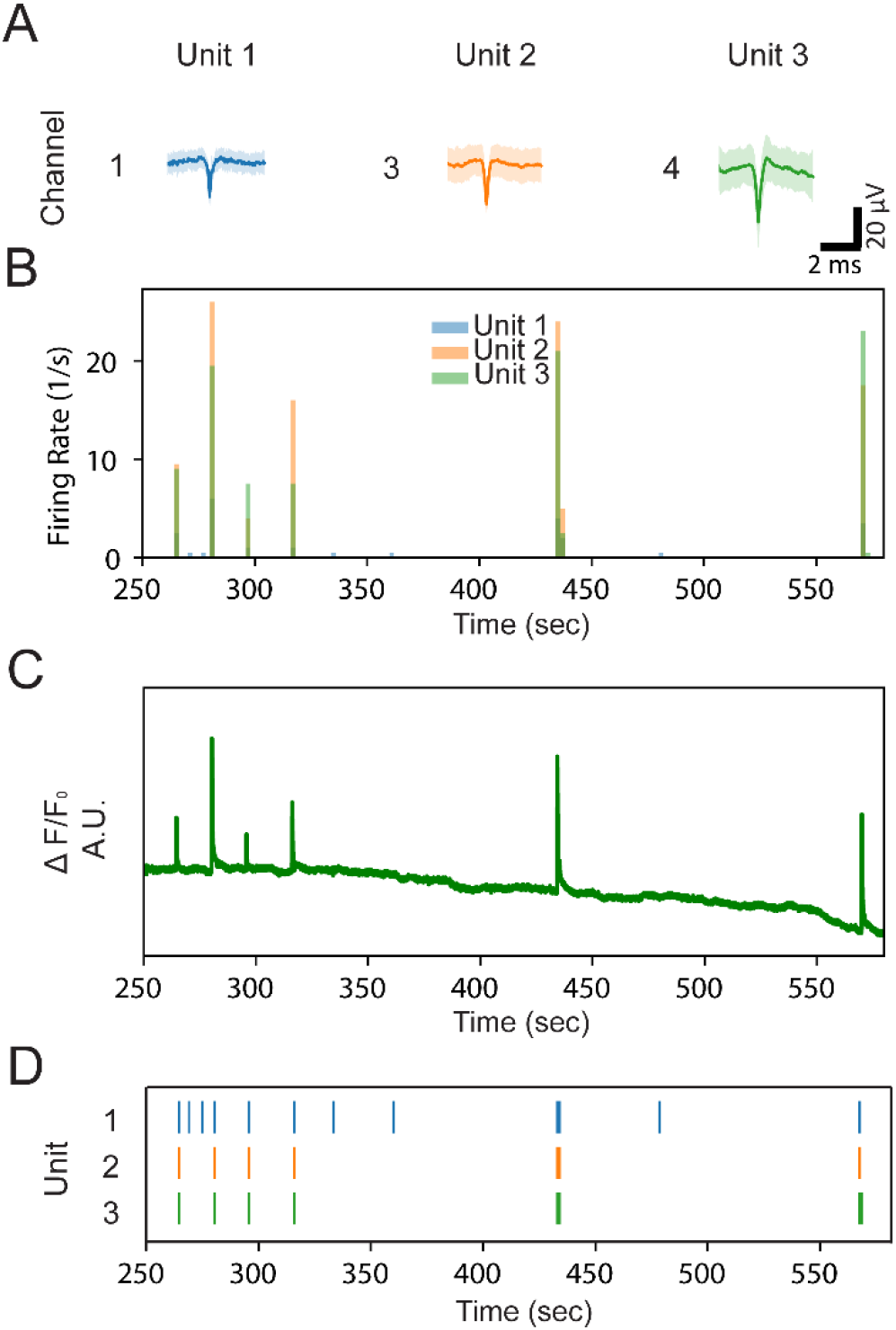
Spontaneous cortical neuron activity correlated to Ca^2+^ fluorescence transients for the second spheroid. **(A)** Averaged spike (solid trace) and standard deviation (shaded region) detected in each unit. The channel where the unit was detected is indicated by the number on the left of the trace. **(B)** Firing rate of representative units, overlaid. **(C)** Optically recorded Ca^2+^ fluorescence intensity from the same time range. **(D)** Raster plots showing firing patterns of detected units over the same time range.

**Figure S12.**
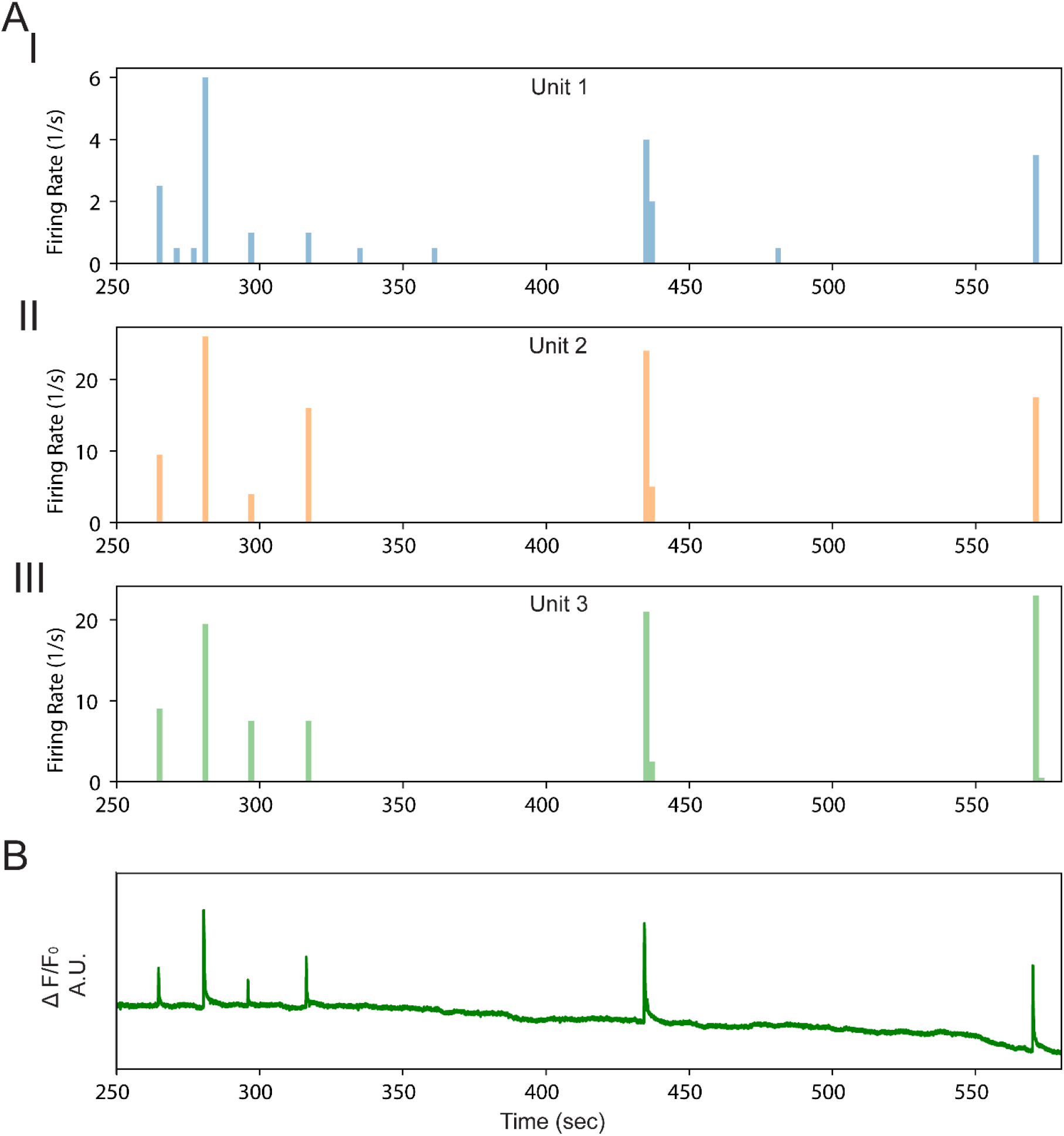
Spontaneous firing of representative units detected after spike sorting of the field potential recording and corresponding whole-spheroid Ca^2+^ fluorescence for second spheroid. **(A)** Firing rate of individual units: (**I**) unit 1, (**II**) unit 2, (**III**) unit 3. **(B)** Optically recorded Ca^2+^ fluorescence intensity from the same time range.

**Figure S13.**
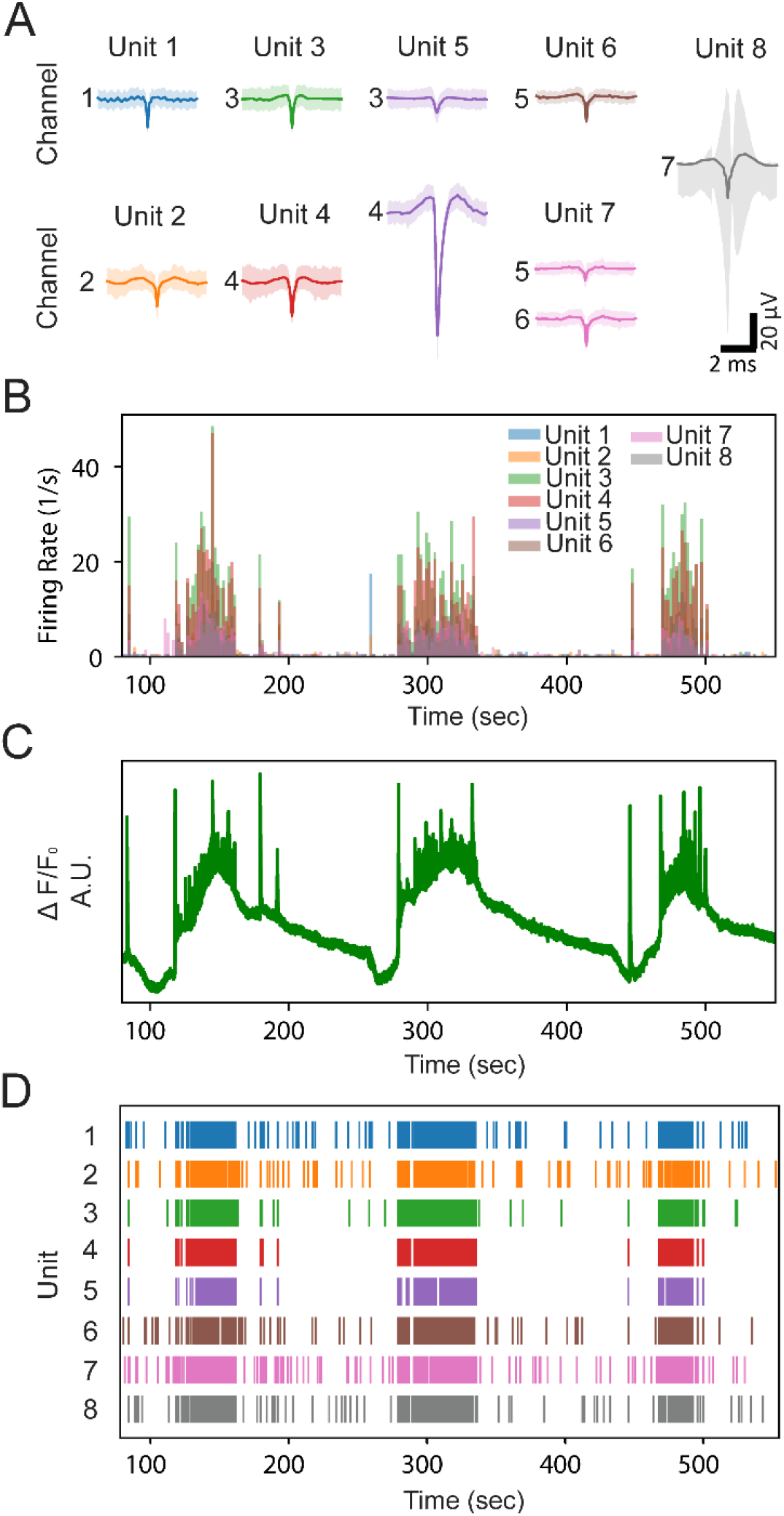
Glutamate-induced cortical neuron activity correlated to Ca^2+^ fluorescence transients for the second spheroid. **(A)** Averaged spike (solid trace) and standard deviation (shaded region) detected in each unit. The channel where the unit was detected is indicated by the number on the left of the trace. **(B)** Firing rate of representative units, overlaid. **(C)** Optically recorded Ca^2+^ fluorescence intensity from the same time range. **(D)** Raster plots showing firing patterns of detected units over the same time range.

**Figure S14.**
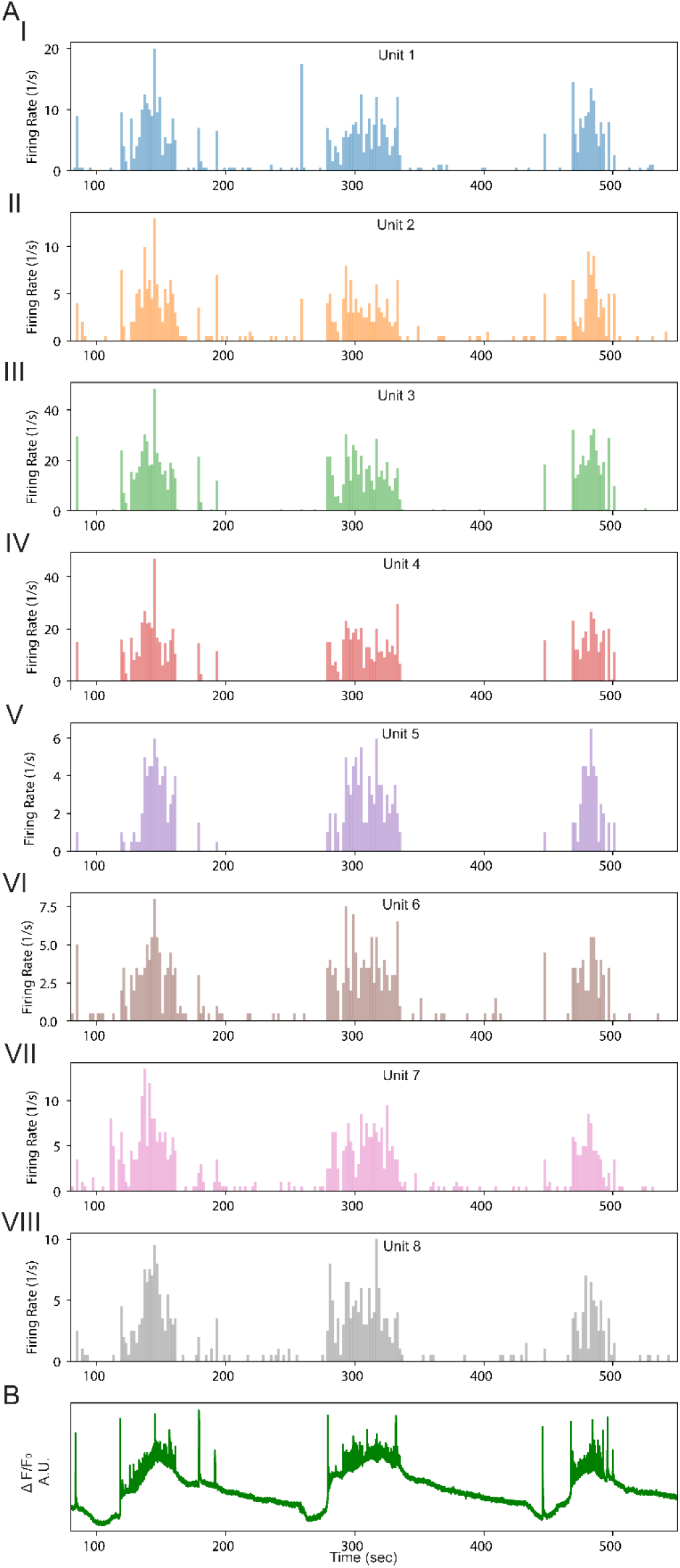
Glutamate-induced firing of units detected after spike sorting of the field potential recording and corresponding whole-spheroid Ca^2+^ fluorescence for second spheroid. **(A)** Firing rate of individual units: (**I**) unit 1, (**II**) unit 2, (**III**) unit 3, (**IV**) unit 4, (**V**) unit5, (**VI**) unit 6, (**VII**) unit 7, and (**VIII**) unit 8. **(B)** Optically recorded Ca^2+^ fluorescence intensity from the same time range.

**Figure S15.**
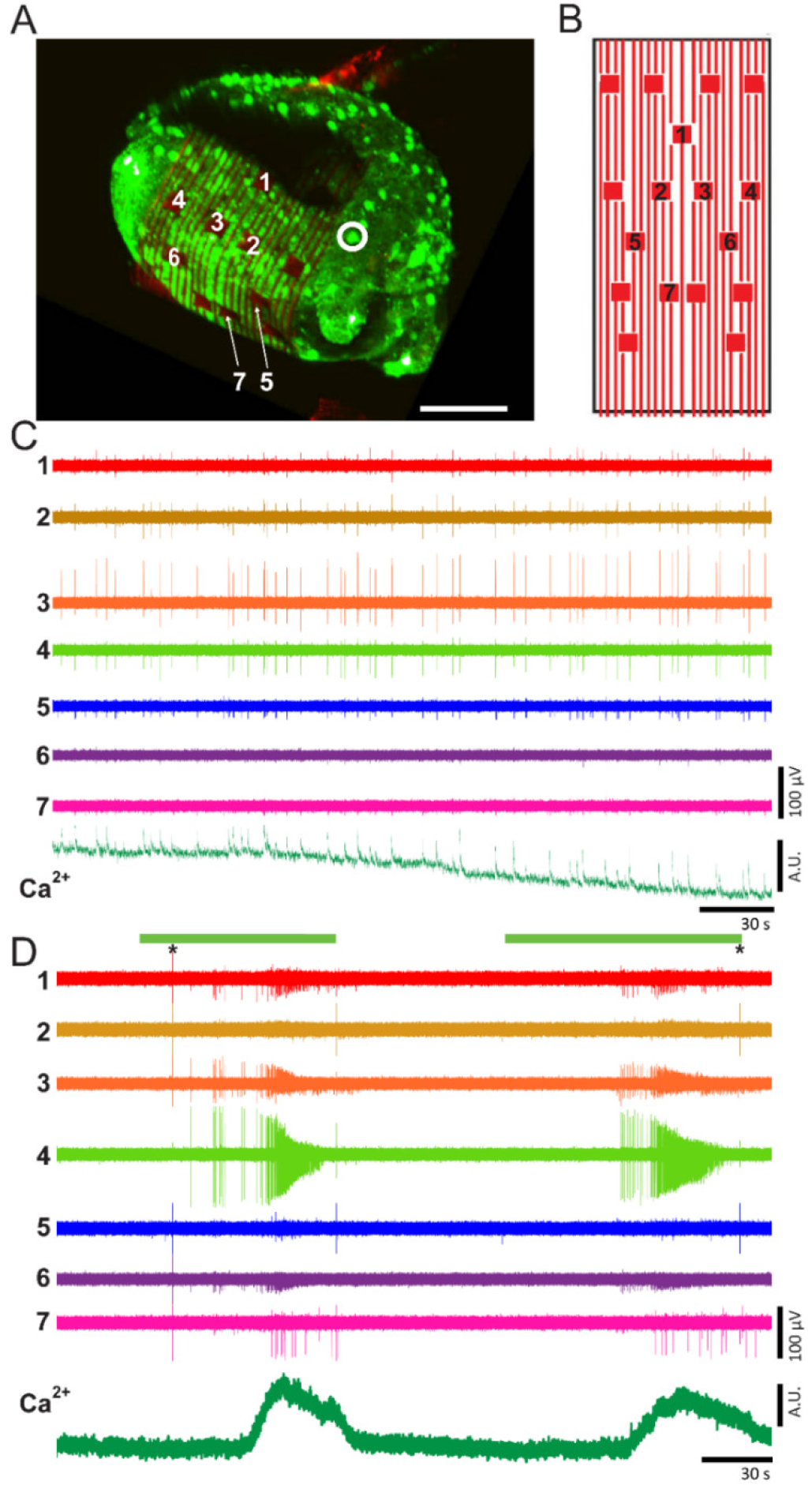
Electrical and optical recordings in 3D of cortical spheroids (third spheroid) using 3D-self-rolled biosensor array (3D-SR-BA). **(A)** A 3D confocal microscopy image of 3D cortical spheroid labeled with Ca^2+^ indicator dye (Cal-520, green fluorescence) encapsulated by the 3D-SR-BA. Scale bar is 100 μm. **(B)** A 2D map of the microelectrodes labeled in panel (A) recording electrical activity of cortical neurons. **(C)** Spontaneously firing neurons recorded via the channels labeled in panel (A) and (B) with simultaneous functional Ca^2+^ imaging. **(D)** Voltage traces recorded from cortical neurons firing upon addition of glutamate and simultaneous functional Ca^2+^ imaging, with fluorescence intensity recorded from a region marked by the white circle. Green bars indicate the time of glutamate flow. **\*** Denotes electrical artifacts.

**Figure S16.**
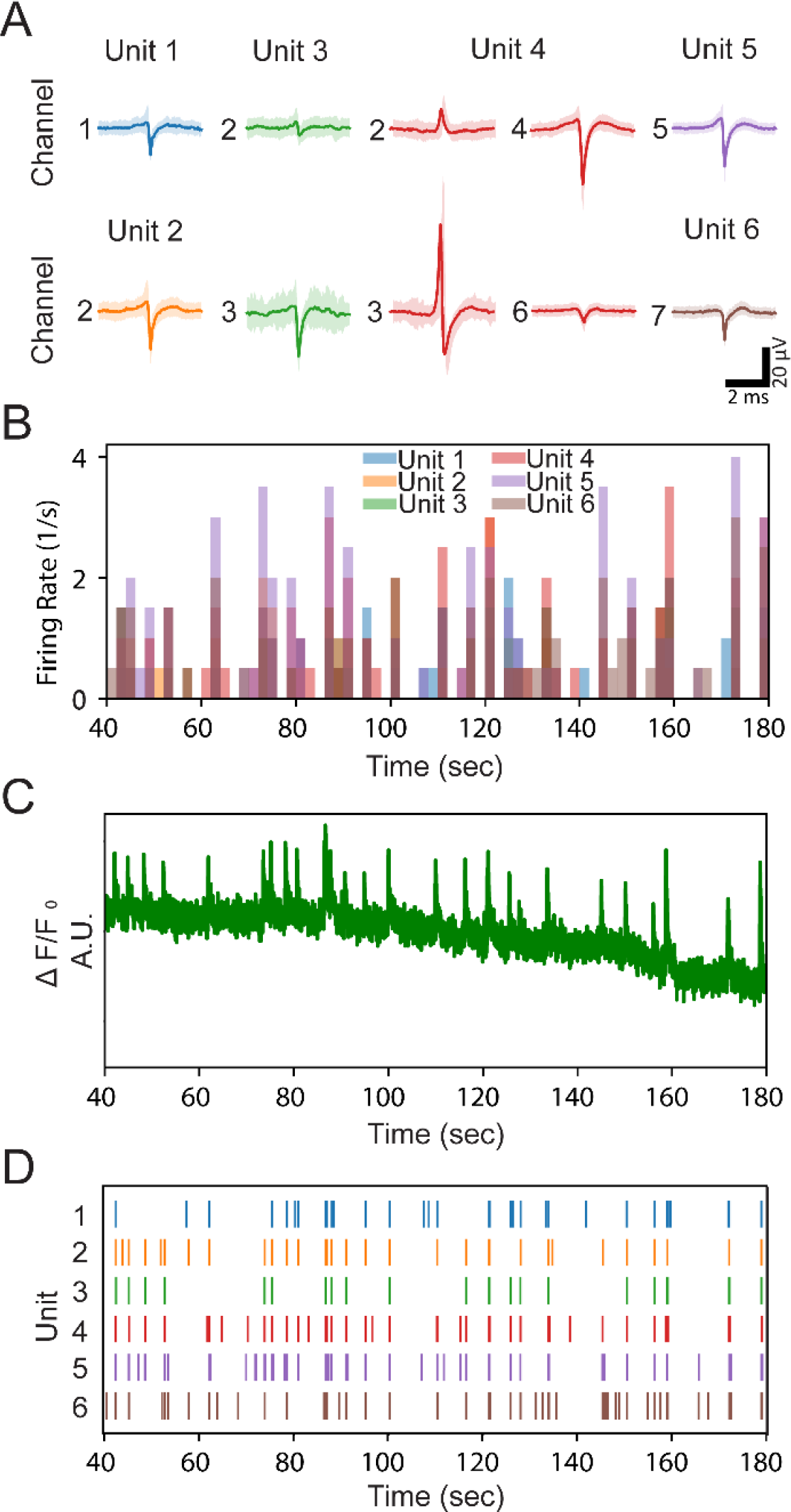
Spontaneous cortical neuron activity correlated to Ca^2+^ fluorescence transients for third spheroid. **(A)** Averaged spike (solid trace) and standard deviation (shaded region) detected in each unit. The channel where the unit was detected is indicated by the number on the left of the trace. **(B)** Firing rate of representative units, overlaid. **(C)** Optically recorded Ca^2+^ fluorescence intensity from the same time range. **(D)** Raster plots showing firing patterns of detected units over the same time range.

**Figure S17.**
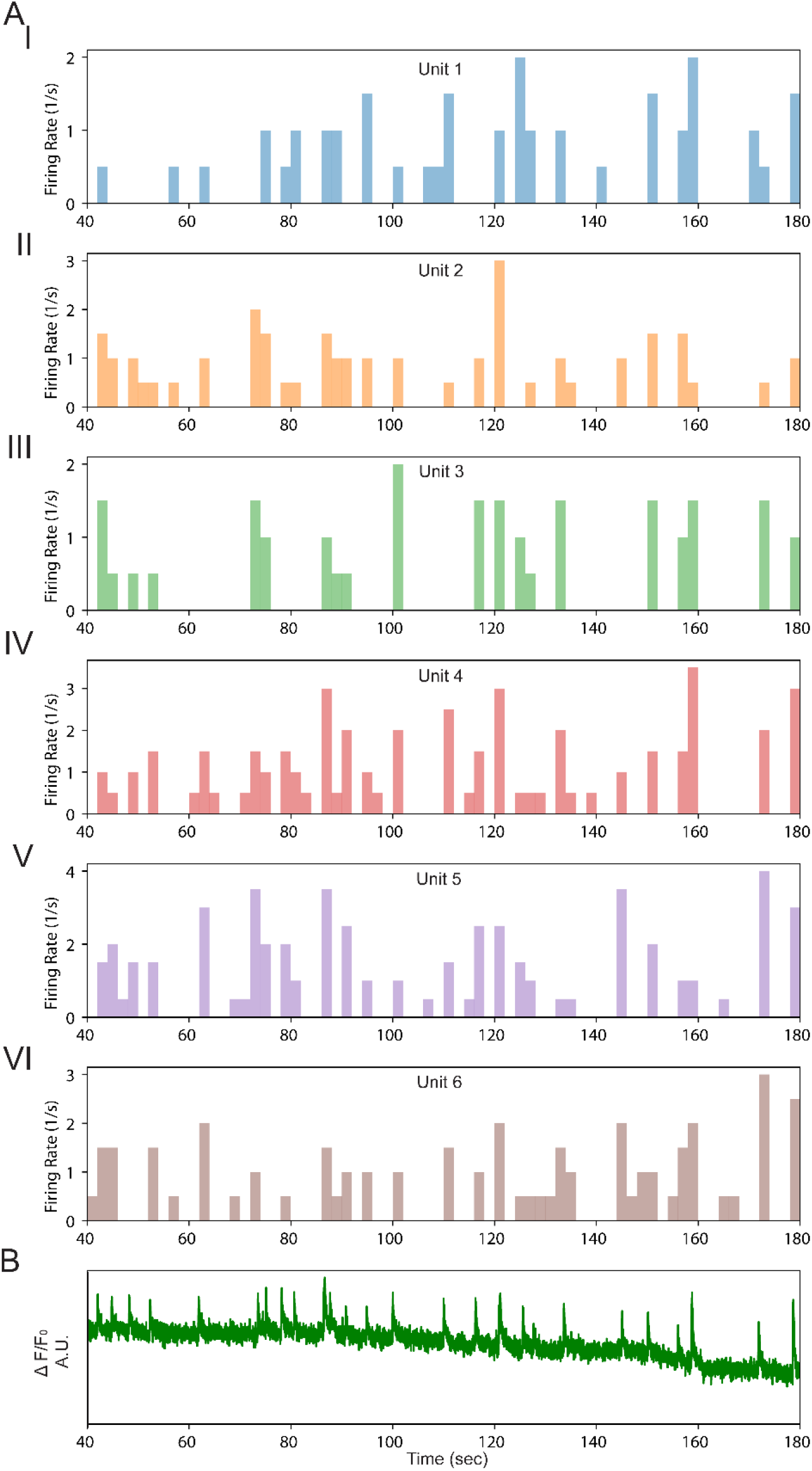
Spontaneous firing of representative units detected after spike sorting of the field potential recording and corresponding whole-spheroid Ca^2+^ fluorescence for third spheroid. **(A)** Firing rate of individual units: (**I**) unit 1, (**II**) unit 2, (**III**) unit 3, (**IV**) unit 4, (**V**) unit 5, and (**VI**) unit 6. **(B)** Optically recorded Ca^2+^ fluorescence intensity from the same time range.

**Figure S18.**
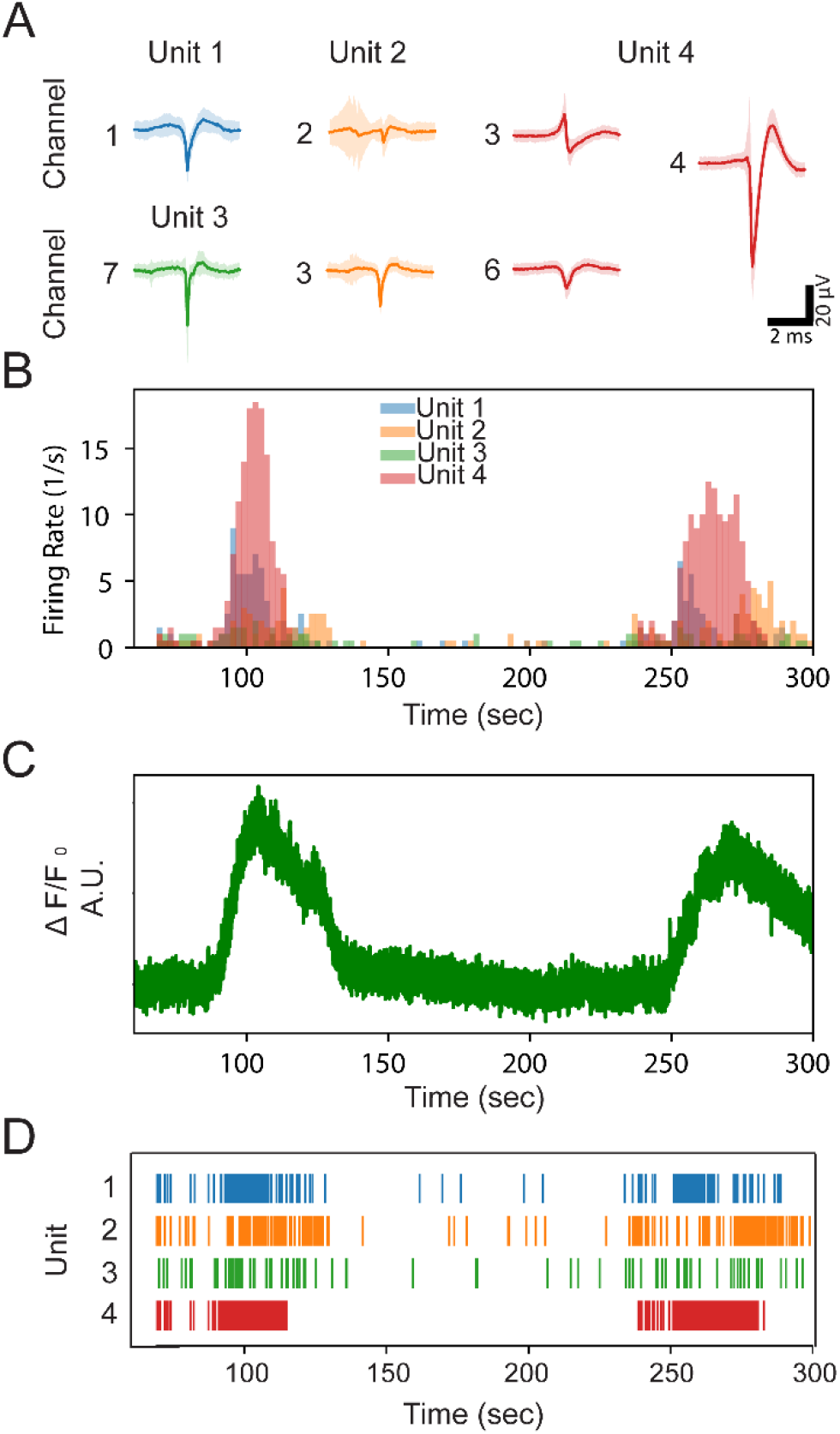
Glutamate-induced cortical neuron activity correlated to Ca^2+^ fluorescence transients for third spheroid. **(A)** Averaged spike (solid trace) and standard deviation (shaded region) detected in each unit. The channel where the unit was detected is indicated by the number on the left of the trace. **(B)** Firing rate of representative units, overlaid. **(C)** Optically recorded Ca^2+^ fluorescence intensity from the same time range. **(D)** Raster plots showing firing patterns of detected units over the same time range.

**Figure S19.**
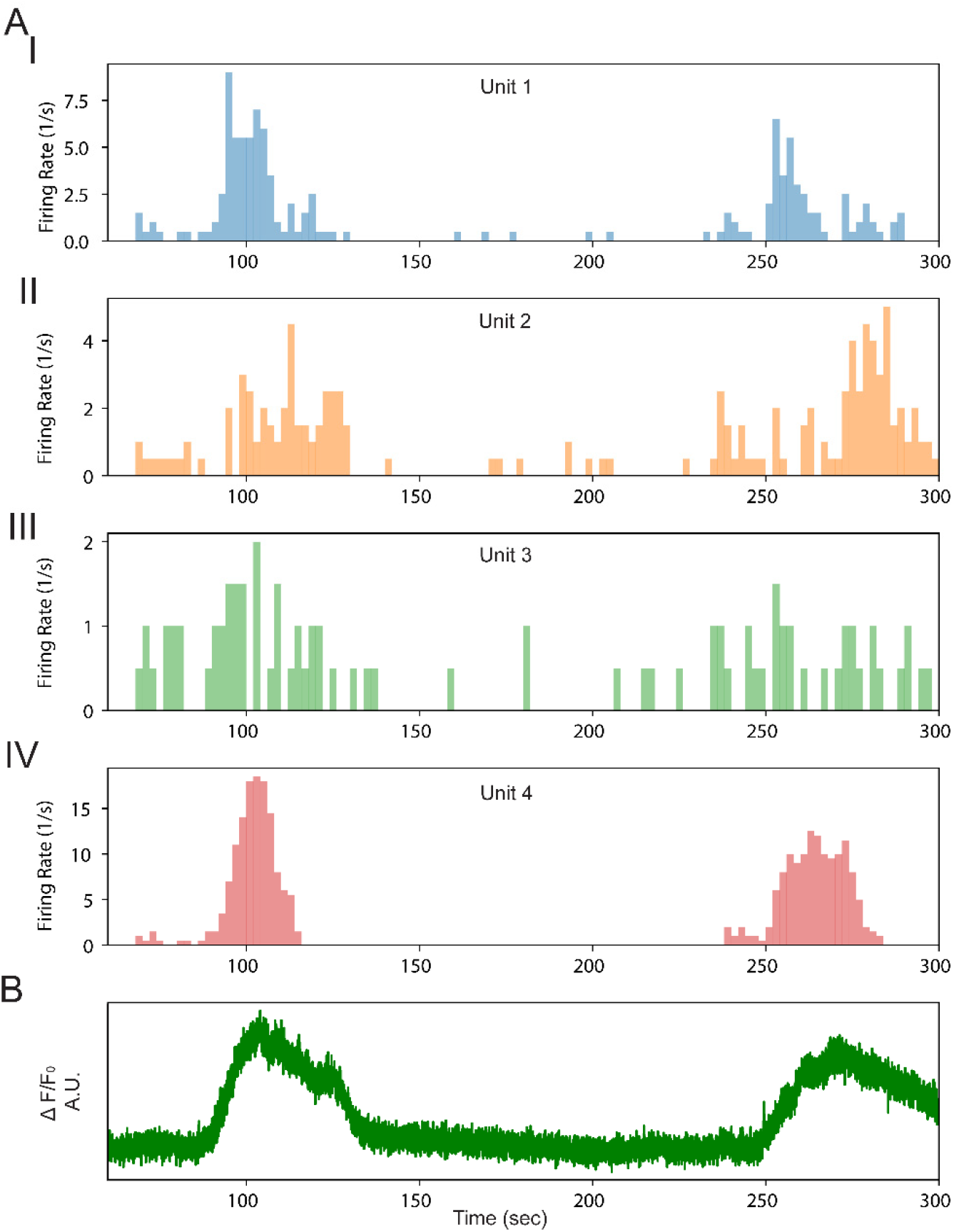
Glutamate-induced firing of representative units detected after spike sorting of the field potential recording and corresponding ROI Ca^2+^ fluorescence for third spheroid. **(A)** Firing rate of individual units: (**I**) unit 1, (**II**) unit 2, (**III**) unit 3, and (**IV**) unit 4. (B) Optically recorded Ca^2+^ fluorescence intensity from the same time range.

